# RNA Editing of DEGS1 Links Genetic Risk to Ceramide Dysregulation and Astrocyte Toxicity in ALS

**DOI:** 10.64898/2026.09.17.752108

**Authors:** Alan Guo, Samuel Lessard, Lilu Guo, Melody Li, Disha Sood, Rachel Passaro, Jeremy Huang, Clement Chatelain, Emanuele de Rinaldis, Shameer Khader, Bailin Zhang, James C. Dodge, Steven Rodriguez

**Author notes:** Correspondence to: Steven Rodriguez; James C. Dodge, Sanofi, 350 Water Street, Cambridge, MA 02141, USA. These authors contributed equally to this work (Alan Guo, Samuel Lessard).

## Abstract

Amyotrophic lateral sclerosis (ALS) is a fatal neurodegenerative disease with ∼90% of cases being sporadic. Although genome-wide association studies have identified numerous genetic risk loci, these variants account for only a portion of ALS heritability, suggesting that additional genetic and regulatory mechanisms contribute to disease risk. Furthermore, the mechanisms linking common variants to disease pathogenesis remain poorly understood. Adenosine-to-inosine (A-to-I) RNA editing is dysregulated in ALS, yet systematic identification of disease-relevant editing events and their functional consequences has been limited.

We performed comprehensive RNA editing analysis across eight central nervous system regions from a Target ALS cohort, identifying 752 differentially edited sites in 304 genes. Global RNA editing was significantly reduced in ALS tissues, particularly in spinal cord and choroid plexus, with altered editing in genes involved in immune signaling, stress response and RNA processing. Through genetic colocalization analysis integrating editing quantitative trait loci (edQTLs) with ALS GWAS data, we identified common variants at the *DEGS1* locus associated with both increased 3′ UTR editing and elevated ALS risk (posterior probability >0.88). *DEGS1* encodes dihydroceramide desaturase 1, a key enzyme converting dihydroceramide to ceramide, the precursor to all sphingolipid species.

Using CRISPR-generated *DEGS1* knockout iPSCs and lipidomic profiling, we demonstrated that DEGS1 is essential for ceramide synthesis and downstream sphingolipid production in astrocytes. Functional validation revealed that ALS-associated editing of the *DEGS1* 3′ UTR significantly increased DEGS1 protein expression. Critically, astrocytes expressing the fully edited *DEGS1* 3′ UTR increased ceramide levels and reduced cell viability compared to those expressing the unedited construct. These findings establish a novel regulatory link between RNA editing, sphingolipid metabolism and ALS pathogenesis. Our results demonstrate that common genetic variants can influence disease risk through post-transcriptional mechanisms that alter lipid homeostasis in astrocytes, a cell type increasingly recognized as central to ALS progression.

## Introduction

Amyotrophic Lateral Sclerosis (ALS) is a fatal neurodegenerative disease that affects the upper and lower motor neurons of the brain and spinal cord, resulting in progressive muscle weakness into paralysis and death within 2-5 years post-diagnosis^1^. Approximately 10% of ALS cases are familial and driven by highly penetrant mutations in genes such as *C9orf72*, *SOD1*, *TARDBP* and *FUS*, whereas 90% are sporadic^2^. Genome-wide association studies (GWAS) have identified ∼15 loci associated with increased risk of ALS^3^; however, these loci account for only a small fraction of the disease heritability estimated from family and twin studies, highlighting substantial missing heritability and suggesting that additional genetic and regulatory mechanisms contribute to disease susceptibility^3–7^. Environmental and epigenetic factors, together with numerous genetic variants of small effect, are also thought to influence ALS risk, age at onset, and disease severity^7–10^. Collectively, these observations suggest that mechanisms beyond conventional genetic variation may contribute to ALS pathogenesis. Yet, because many GWAS risk variants exert modest effects and likely influence diverse disease-relevant traits and pathways, integrative functional analyses are essential to elucidate how these variants contribute to disease pathogenesis.

Adenosine-to-inosine (A-to-I) or sometimes referred to as adenosine-to-guanosine (A-to-G) RNA editing is dysregulated in ALS. For example, Adenosine Deaminase Acting on RNA (ADAR) enzymes are decreased in the spinal cords of sporadic ALS patients^11,12^. Notably, decreased editing efficiency at the Q/R site of GluA2 subunit of AMPA receptors in ADAR2^-/-^ mice leads to increased Ca^2+^ permeability, excitotoxicity and ultimately motor neuron death^11–15^. Separately, RNA editing of dsRNA by ADAR1 suppresses the innate immune pathway, which is also implicated in the disease progression of ALS^16–20^. Furthermore, GWAS enrichment analyses of editing quantitative trait loci (edQTLs) demonstrated that inflammatory disease risk variants can influence A-to-I RNA editing, suggesting that altered RNA editing may represent a broader mechanism linking genetic variation to disease susceptibility, including in ALS^21^. These observations underscore the need for systematic identification of differentially edited transcripts in ALS, with prioritization based on their genetic association with disease risk and potential disease-modifying effects.

Ceramides are lipids that play vital roles in complex sphingolipid and glycosphingolipid synthesis, membrane function and cell signaling. Disrupted ceramide homeostasis is implicated in multiple cell-death pathways and several neurologic diseases^22–25^. Several lines of evidence indicate that ceramide and its derived lipids play a key role in the pathogenesis of ALS. Gain of function mutations in SPTLC1/2, encoding the rate limiting enzyme for *de novo* ceramide synthesis, lead to excessive sphingolipid synthesis and cause juvenile-onset ALS^26,27^. Genetic evidence further supports a pathogenic role for ceramide accumulation in motor neuron degeneration, as partial loss-of-function mutations in ASAH1, the lysosomal enzyme that degrades ceramide, cause early-onset spinal muscular atrophy^28,29^. While findings from juvenile-onset ALS and related motor neuron disorders support a pathogenic role for ceramide dysregulation, direct genetic links between sphingolipid pathway genes and the more common familial and sporadic forms of ALS remain limited. ALS patients and mouse models exhibit consistent and progressive disruptions in ceramide and sphingolipid metabolism. Targeted lipidomic profiling of postmortem spinal cord tissue demonstrated accumulation of ceramide and its metabolic derivatives, including sphingomyelin (SM) and glycosphingolipids, consistent with widespread perturbations in lipid metabolism in ALS^30–33^. Longitudinal plasma lipidomic analyses showed progressive increases in short- and very long-chain ceramide species together with reductions in SM species and the rate of SM depletion tracked disease progression as measured by ALSFRS-R^34,35^. In CSF, researchers also detected elevated SM and glucosylceramide levels^36^. Together, these findings establish dysregulated sphingolipid flux as a contributor to ALS pathogenesis and a biomarker of disease progression.

Here, we identify common ALS associated genomic variants linked to RNA editing of *DEGS1*, providing evidence of a genetic connection between ceramide metabolism and sporadic adult-onset ALS. We show that global RNA editing is reduced in ALS spinal cords and identify multiple differentially edited genes. Through colocalization analysis, we identify the ceramide-synthesizing enzyme DEGS1 as exhibiting increased RNA editing in ALS and investigate the functional consequences of ALS-associated editing sites within its 3′ UTR. We identify ALS-associated editing events within the *DEGS1* 3′ UTR and show that DEGS1 is essential for sphingolipid synthesis in astrocytes. Importantly, 3′ UTR editing enhances DEGS1 protein expression, leading to increased ceramide levels and reduced astrocyte viability. Together, these findings reveal a previously unrecognized regulatory link between RNA editing, sphingolipid metabolism and ALS pathogenesis, with enrichment of ALS GWAS signals at the *DEGS1* locus further supporting its contribution to disease susceptibility.

## Materials and methods

### ALS Cohort and data

Whole-genome sequencing (WGS) and bulk RNA-sequencing data were obtained from the Target ALS post-mortem cohort (NYGC ALS Consortium; 2020 and 2022 freezes)^37,38^. RNA-seq was retained for eight central-nervous-system regions: cerebellum, choroid plexus, frontal cortex, lateral and medial motor cortex and cervical, lumbar and thoracic spinal cord.

### WGS quality control

We processed Joint-called GATK VCFs^39^ with the WGS-QC pipeline^37,40^ on genome build hg38. Briefly, the pipeline removed ENCODE blacklist regions^41^, retained biallelic GATK-PASS variants and applied genotype-level read depth (DP ≥ 10), genotype quality (GQ ≥ 20), SNP missingness (< 0.15), mapping quality (58.75 ≤ MQ ≤ 61.25), VQSLOD (≥ 7.81; SNPs only), an inbreeding-coefficient filter (Inbreeding Coef ≥ -0.8), per-sample missingness (< 0.1) and a KING relatedness threshold (≤ 0.125). QC used bcftools^42^ 1.9, vcftools^43^ 0.1.15, KING^44^ 2.1.6, vcflib^45^ and PLINK2^46^. We applied an additional minor allele frequency filter (MAF > 0.05) for downstream eQTL/edQTL mapping. We further verified genetic sex against reported sex. For ancestry, QC’d genotypes were merged with LD-pruned 1000 Genomes phase-3 reference samples^47^ and projected by principal component analysis (PCA). Samples were assigned to super-populations and the analysis was restricted to genetically European participants, defined as Target ALS samples within ±4 SD of the 1000G European mean on PC1 and PC2.

### RNA-seq sample QC

QIAGEN Digital Insights Professional Services processed raw FASTQ files in OmicSoft Suite 11.7.3.3 (QIAGEN, Redwood City, CA) using a standardized RNA-seq ingestion pipeline with default settings. In brief, the pipeline performed read and sample quality control, alignment to the Genome Reference Consortium Human Build 38 (GRCh38) with OSA4^48^ gene-level expression quantification and FPKM normalization using an expectation-maximization algorithm^49^ with the OmicsoftGenCode.V33 (GENCODE 33) gene model and differential expression analysis with DESeq2 1.10.14 comparing tissue-specific ALS patients to non-neurological controls. We excluded ALS patients with multiple neurological conditions from pairwise differential expression analysis and we considered differentially expressed genes (DEGs) as significant with a false discovery rate (FDR) adjusted *p* < 0.05. Sample identity (meta-data) and tissue assignment were verified by PCA (top 10 PCs) and UMAP of the FPKM matrix. A sequencing-platform batch effect was identified and a small number of mislabeled cerebellum and choroid samples were removed (flagged as PCA outliers). Biological sex was further confirmed by evaluating the expression of Y- and X-specific genes (*RPS4Y1*/*XIST*). For subjects with multiple samples of the same tissue, the sample with the highest RIN was retained.

### RNA-editing quantification

We performed RNA-editing quantification^50^ and edQTL mapping following the GTEx edQTL pipeline^21^ with the published code at (https://github.com/vargasliqin/mpileup; https://github.com/vargasliqin/GTEx_edQTL) adapted to the Target ALS cohort. Briefly, editing was quantified against a reference A-to-I site list, which combined known sites from the RADAR database^51^, tissue-specific sites identified in GTEx V6p and published hyper-editing sites^51,52^ and containing over 2,802,002 sites. For each BAM, reads were counted at each catalogued site using samtools mpileup^42^ with minimum base quality 20, minimum mapping quality 5 and option “-A -B -p 1000000”. The strand-aware editing level was computed as the fraction of edited over total reads: G/(A+G) on the plus strand and C/(C+T) on the minus strand^21^. We restricted the quantification to uniquely mapped reads (‘NH:i:1’) and retaining autosomal sites covered by ≥ 20 reads in ≥ 20 samples per tissue with non-zero variance across samples. Finally, we performed rank-based inverse-normal quantile normalization of editing ratios across samples within each tissue. This yielded between 29,046 and 44,480 testable editing sites per tissue (Supplementary Table 1).

### Differential editing

For each tissue and editing site, the quantile-normalized editing level was modeled by linear regression as a function of disease status (ALS vs non-neurological control), adjusting for sex and sequencing platform. Benjamini–Hochberg false discovery rate (FDR) was used for multiple testing correction, with FDR < 0.05 treated as significant. Sites were annotated to genes via their overlap with gene bodies based on the R package TxDb.Hsapiens.UCSC.hg38.knownGene. Sites overlapping with more than one gene are reported against each. Skew of differential editing directionality was assessed using a binomial sign test included all sites with *P*<0.05. For full list of differentially edited genes and individual edit sites refer to Supplementary File 1.

Genes harboring differentially edited sites were tested for pathway enrichment using QIAGEN Ingenuity Pathway Analysis (IPA)^53^. For each tissue, the differential editing results were reduced to one site per gene based on the site with the lowest nominal p-value. Genes were included in the IPA core analysis based on an absolute effect estimate > 0.5 and nominal p < 0.001, with genes having at least one quantified edited site as background. We report enriched canonical pathways with p-value < 0.05. For full list pathways affect see Supplementary File 2.

### cis-eQTL and cis-edQTL mapping

QTLs were mapped per tissue using tensorQTL^54^ (cis_nominal mode, ±1 Mb cis-window from the editing sites or transcription start sites, MAF ≥ 0.05). Latent factors were estimated with MOFA2^55^. For eQTL input, FPKM values were log-transformed (log(FPKM+1)) and the platform effect was removed using limma removeBatchEffect^56^. Covariates comprised the top 5 genotype PCs, age, genetic sex, sequencing platform (edQTL only) and the top 10 latent expression or editing factors.

### Fine-mapping and colocalization with ALS GWAS

ALS GWAS^3^ summary statistics (GCST90027164; 27,205 cases and 110,881 controls) were munged (MungeSumstats^57^) and lifted to hg38 (UCSC liftOver^58^) for alignment with the QTL coordinates. Independent loci with suggestive association (*P* < 5x10^-6^) were identified by clumping the GWAS summary statistics (PLINK^46^; r^2^< 0.1, 1000G European linkage disequilibrium (LD) reference, 1 Mb window). For each suggestive GWAS locus, all FDR-significant Target ALS edQTLs (q < 0.05) in the region were tested for colocalization with the GWAS signal using coloc^59^. Loci with posterior probability of a shared causal variant (PP.H4) > 0.8 were considered to colocalize. For each candidate locus (PP.H4 > 0.8), both the GWAS and the Target ALS edQTL were fine-mapped independently with SuSiE^60,61^ using a 1000G European LD reference for the GWAS in-sample LD for edQTL loci, computed from the Target ALS WGS genotypes (PLINK^46^ --r square). SuSiE credible sets were derived from the respective summary statistics and LD matrices. Where both traits converged to credible sets, the shared-variant posterior was recomputed with SuSiE-coloc^62^ (per-variant SNP-level PP.H4).

### Comparison with GTEx edQTLs and eQTLs

For each identified ALS-edQTL shared loci, we also evaluated colocalization of independent eQTL^63^ and edQTL^21^ from related GTEx tissues (v8 spinal-cord cervical edQTLs). GTEx eQTL were retrieved from the eQTL Catalogue (QTD000201^64^). Target ALS eQTLs at the same loci were also tested to assess whether the editing signal is accompanied by an expression signal.

### Dual Luciferase vector cloning and assay

3’ UTR of edited or unedited human *DEGS1* was synthesized and cloned by GENEWIZ into pmirGLO vector (Promega) using NheI and SalI restriction enzymes. For edited versions of the 3’ UTR, specific adenosines that corresponded to those found in our bioinformatics pipeline were changed to guanosines. For information on the sequence of the edited 3’ UTRs refer to Supplementary File 3. Dual luciferase constructs were transfected into HeLa using Lipofectamine 2000 (Invitrogen) or human astrocytes using Lipofectamine LTX (Invitrogen). One day post-transfection, Dual-Glo Luciferase assay (Promega) was performed following the manufacturer protocol in 96-well plate format. Luminesce was measured using plate reader.

### Generation of DEGS1 KO iPSCs

To generate cas9-inducible iPSCs, Wildtype PGP1 iPSCs were infected with a lentivirus containing a dox-inducible Cas9 cassette containing blasticidin resistance marker (Horizon Discovery) using an MOI of 1. Successfully transduced cells were selected with blasticidin and clonal selection was performed using serial dilution to generate individual clones. To generate DEGS1 KO iPSCs two lentiviral dox-inducible gRNAs (Cellecta, pRSGTEP-U6Tet-sg-EF1-TetRep-2A-Puro) targeting two different regions of exon 1 of *DEGS1* were pooled and transduced into cas9-inducible iPSCs using an MOI of 2.5. *DEGS1* gRNA sequences are GGTATCACATGGATCATCAT and CGAAGTCTTCCCGCGAGACG. The lentiviral gRNAs used contained a puromycin resistance marker which was used to isolate successfully transduced cells. Clonal selection was performed using serial dilution to generate individual clones. Successful *DEGS1* knockout was confirmed using western blot and targeted lipidomics.

### iPSC differentiation into astrocytes

Astrocyte differentiation protocol was adapted from Rapino et al. 2023^65^. Briefly, iPSCs were maintained and expanded in mTeSR Plus Basal Medium. Upon reaching confluency, iPSCs were dissociated into single cells with Accutase and counted. iPSCs were seeded into 125 mL spinner flasks containing 100 mL of mTeSR medium at a density of 1 × 10⁶ cells/mL, supplemented with 10 μM ROCK inhibitor Y-27632. After 48 hours, the medium was replaced with KSR medium containing 10 μM SB431542 and 1 μM dorsomorphin. For the first five days, the media was refreshed daily. From day 6 to day 12, the media was changed every other day using Neurobasal medium supplemented with dorsomorphin and various cytokines as specified: Day 6 and 8, FGF2 (10 ng/ml) and EGF (10 ng/ml); Day 10 and 12, FGF2 (20 ng/ml), EGF (20 ng/ml) and CNTF (20 ng/ml); Day 16 onward, medium changes continued every two days using Neurobasal medium with 2X N2 supplement and CNTF (20 ng/mL). On day 30, the resulting spheres were dissociated and either cryopreserved or expanded on poly-L-lysine (PLL)-coated plates and grown in Astrocyte media (ScienCell).

### Lipid profiling using liquid chromatography/mass spectrometry

For iPSCs, cells were grown in mTeSR plus media. d17-sphinganine stock was made by adding 200 proof ethanol, sonicated for 5 mins in 37C water bath for a stock concentration of 2mM. d17-sphinganine was sonicated for 20 mins at 37C and added to cell media for at a final concentration of 1uM. For baseline conditions, ethanol was diluted in PBS. After 2 or 24 hours, cells were washed with PBS and flash frozen for lipid profiling. For iPSC-astrocytes, cells were grown in astrocyte media (ScienCell). 3 days before collection, media was replaced with astrocyte media without FBS. Cells were washed with PBS prior to collection. Trypsin (0.025%) was added to each well and incubated for 8 minutes. Cell suspension was collected followed by automated cell counting. Cells were pelleted, supernatant was removed and cell pellet was flash frozen on dry ice for lipid profiling. Data readouts using iPSC-astrocytes were normalized total cell counts.

Cell pellet was extracted with 0.2 mL of extraction solution containing internal standard (methanol/acetonitrile, 1:1 containing 10ng/mL C16d31-Cer, 20ng/mL d35-C18GalCer, 10ng/mL d7-Sphingosine and 10 ng/mL d7-sphinganine, as internal standards). The mixture was vortexed for 10 minutes in multi-tube vortex, sit on ice for 10min and vortex again for 10 min. The homogenate was spined down for 10 minutes at 13,000 g. The supernatant (∼200 μL) was carefully transferred to a MS vial for analysis.

Standard curves (0.01-500 ng/mL) were prepared in the same way as samples with corresponding standards from each lipid class: Ceramide_C16, dH-Ceramide_C16, Sphingosine, Sphinganine, GluCer(d18:1/18:0) GalCer (d18:1/18:0).

LC-MS/MS analysis was conducted on a Waters Acquity UPLC system coupled with a Sciex QTRAP6500 mass spectrometer using multiple-reaction monitoring (MRM). Data were acquired and analyzed by MultiQuant 3.0 (AB Sciex). Calibration curves were constructed by plotting the corresponding peak area ratios of analyte/internal standard versus the corresponding analyte concentrations using 1/x weighing linear regression analysis.

Cer and dHCer (including both d17-Sphingosine backbone from labeled substrate and the endogenous d18-backbone ceramides), the Ceramide profile is separated by a Waters Acquity UPLC BEH C8 (100 × 2.1 mm, 1.7um particles, cat#186002878) at a flow rate of 0.4 mL/min. The column is heated at 60 C during the run. The mobile phase consists of A) 0.2% Formic Acid; 5 mM Ammonium formate in DI water and B) 0.2% Formic Acid; 5 mM Ammonium formate in (50:50) Methanol:Acetonitrile. The injection volume was 5 μL and the total runtime was 6 min. The step gradient was as follows: 0–0.2 min, 85% solvent B; 0.2–1.8 min, 85 to 98% solvent B; 1.81-3.4 min, 100% solvent B, 3.4–3.5 min, 100 to 85% solvent B; 3.5–6 min 85% solvent B. The ESI+ source temperature was 450 °C; the ESI needle was 4,500 V; the declustering potential was 60 V; the entrance potential was 10 V; and the collision cell exit potential was 10 V. The collision and curtain gas were set at medium and 15, respectively. GS1 and GS2 were set at 80 and 20. The collision energy was 34 eV for Ceramide and 40 eV for dHCer. For MRM, the dwell time was set at 50 ms for each transition.
GluCer and GalCer (including both d17-Sphingosine backbone from labeled substrate and the endogenous d18-backbone glucosylceramides and galactosylceramides) Analyte (1 μl) was injected into the system with a partial loop in the needle overfill mode. GluCer and GalCer (stereoisomers) are separated on a waters Cortecs HILIC 2.1 × 100 mm 2.7 μm particles (Waters, 186007427). The mobile phase consisted of Solvent A: 5 mM Ammonium Acetate in 96:2:1:1 (v/v/v/v) of Acetonitrile:Methanol:Acetic Acid:Water and Solvent B: 5 mM Ammonium Acetate in 80:20:1 (v/v/v) Methanol: Water:Acetic Acid. The gradient program was isocratic 98% A at 0. 5 mL/min for 5 minutes per sample. An ESI+ source was used with the following parameters: curtain gas 20.0, ionSpray voltage, 4.5 kV; temperature, 450 °C; Ion source gas: 80 and 80, Declustering potential 120, entrance potential 10, collision energy 47 and collision cell exit potential: 10.
Sphinganine (Sa) and Sphingosine (So) (including both d17- and d18-backbone) So and Sa are analyzed by a Waters Acquity UPLC BEH C18 2.1x50mm 1.7um, (186002350) at a flow rate of 0.5 mL/min. The mobile phase consists of A) Water with 0.1% Formic acid and B) 85:15 Methanol/Acetonitrile with 0.1% formic acid.

The injection volume was 5 μL and the total runtime was 5 min. The step gradient was as follows: 0–0.2 min, 50% solvent B; 0.2–3 min, 50 to 99% solvent B; 3-4 min, 99% solvent B, 4–4.1 min, 99 to 50% solvent B; 4.1–5 min 50% solvent B.

The ESI+ source temperature was 350 °C; the ESI needle was 5,500 V; the declustering potential was 60 V; the entrance potential was 40 V; and the collision cell exit potential was 10 V. The collision and curtain gas were set at medium and 10, respectively. GS1 and GS2 were set at 40 and 40. The collision energy was 15 eV.

### Western Blotting

Media was removed from wells and cells were washed with ice-cold PBS. PBS was removed and RIPA buffer containing protease/phosphatase inhibitor cocktail was added directly to the wells, pipetted up and down and incubated on ice with occasional shaking for 10 minutes. Cell content was collected, vortexed for one minute and centrifuged to separate debris from cell lysates. Protein concentration was determined using Pierce BCA protein assay and protein concentration was normalized across all samples for each run. Sample buffer and reducing agent were added to protein lysates and boiled at 95C for 10 mins prior to loading on NuPAGE gel and run for 35 mins at 200V. Gel was removed and subjected to wet transfer using PVDF membrane for 1 hour at 30V. The membrane was incubated in Licor blocking buffer for 1 hour, incubated in primary antibodies in Licor antibody dilution solution overnight at 4C and washed 3 times at 5 minutes with TBST. DEGS1 (Abcam, ab185237) and β-Actin (Cell Signaling Technologies, 3700S). TBST was removed and membranes were incubated in secondary antibody in Licor antibody dilution solution for 2 hours before imaging in Licor Odessey.

### Immunocytochemistry and imaging

Cells were plated in 96 well PhenoPlate (Revvity) to confluency. Media was removed from wells and cells were gently washed with PBS. Cells were fixed using 4% PFA in PBS for 10 mins and gently washed with PBS. The following steps apply to all conditions except for cells stained with CD44 in which Triton-X was removed for all solutions. Cells were permeabilized in 0.2% Triton-X in PBS for 10 mins. Permeabilization solution was removed and cells were incubated in blocking buffer (5% BSA, 0.1% Triton-X in PBS) for 1 hour at room temperature in the dark. Blocking buffer was removed and cells were incubated with primary antibodies in blocking buffer for overnight at 4°C. The next day, primary antibody solution was removed and cells were gently washed with PBS 3 times for 5 minutes each. Cells were then incubated in secondary antibody and Hoechst in blocking buffer for 2 hours at room temperature in the dark. Secondary antibody solution was removed and washed PBS 3 times for 5 minutes each. Plates were imaged using Opera Phenix Plus. CD44 is from abcam (ab157107), S100β is from abcam (ab52642).

### Cell viability assay

100k DEGS1 KO iPSC-astrocytes were electroporated using Amaxa 4D nucleofector in 16-strip format using the following conditions below: For all conditions, cells were electroporated using CA137 program and cultured in Astrocyte media without FBS (ScienCell). Cells were transfected with DEGS1 with 1ug WT 3’ UTR, DEGS1 edALL 3’ UTR or 0.5ug GFP (molar equivalent). After transfection, 200ul of media was added to each of the wells and plated in 96-well format and media was changed the next day. Cells were maintained for 10 days with media changes every 3-4 days. On day 10, 100ul of cell media was removed and 100ul of CellTiter-Glo 2.0 reagent (Promega) was added directly to each well, placed on shaker for 2 mins, 300 RPM, incubated in the dark for 10 mins and luminescence was measured using plate reader.

### 3’ UTR and miRNA mapping

miRNA to 3’ UTR mapping prediction mapping was performed using AI coding platform, Cursor. Predicted miRNA binding to either unedited (WT) or edited (ed1 and edALL) human *DEGS1* 3′ UTR sequences was assessed using TargetScan-style seed rules applied to the mature miRNA sequence (positions 2–8 for 7mer-m8 and 8mer; positions 2–7 plus an adenine opposite miRNA position 1 for 7mer-1a; 8mer additionally requiring an adenine opposite position 1 relative to the 7mer-m8 core). Mature human miRNAs were taken from the miRBase mature sequence compendium. For each UTR, each mature miRNA was classified independently for 8mer, 7mer-m8 and 7mer-1a using exact matching of the corresponding DNA reverse-complement patterns using Biostrings in R. A miRNA was counted as present if it matched to a given seed type. Summary bar plots were generated with ggplot2.

For cross-referencing unique miRNAs whose binding sites were gain or lost due to RNA editing to existing ALS literature, associations with ALS were taken from HMDD v4.0^66^ with text matching ALS-related terms. miRNA names were normalized and matched to HMDD gene-level IDs. PubMed was queried per miRNA (NCBI E-utilities via rentrez) with ALS-related terms in the title or abstract. We recorded HMDD yes/no, PubMed hit counts, sample PMIDs and which seed types (8mer, 7mer-m8, 7mer-1a) the miRNA had in the gain/loss set. HMDD and PubMed are association/co-occurrence checks only and not mechanistic or DEGS1-specific evidence. For full summary of miRNA binding site changes see Supplementary File 4.

## Results

### RNA editing is altered in ALS spinal cords

To understand whether RNA editing is dysregulated in ALS, we leveraged a Target ALS dataset^37,38^, including transcriptomic samples across eight central-nervous-system regions, with sample sizes ranging from 59 to 202 (Fig. 1A, Supplementary Table 1). We quantified A-to-I (A-to-G) RNA editing across all transcripts and performed differential editing analysis between ALS vs. non-neurological controls to identify sites dysregulated in the disease. We detected significant differential editing (False discovery rate < 0.05) in five of the eight regions (Fig. 1B). Overall, we identified 752 differentially edited sites across 304 genes overall. The signal concentrated in cervical spinal cord (352 sites / 151 genes), lumbar spinal cord (284 sites / 138 genes) and choroid plexus (109 sites / 87 genes) – sites strongly affected by ALS. In contrast, RNA editing alterations were sparse in the cerebellum and medial motor cortex, with only three and four differentially edited sites detected, respectively. Outside the cortex, we observed a global trend where RNA editing was lower in ALS compared to controls (binomial *P*<1x10^-4^, Fig. 1C, Supplementary File 1), particularly pronounced in choroid (*P*=6x10^-34^). Some of the strongest dysregulated sites included increased editing of *SYT11* (synaptotagmin-11) in spinal cord (*P* = 8×10⁻¹²), together with altered editing of the microglial and lysosomal-related genes *CD68*, *CTSS* and *CTSB*, the long non-coding RNA *NEAT1*, the splicing factors *U2AF1* and *SNRPD3* and the double-stranded-RNA sensor *EIF2AK2* and the type-I interferon receptor subunits *IFNAR1* and *IFNAR2*. These findings align with the broader observation that A-to-I editing is dysregulated in ALS motor neurons, where loss of ADAR2 activity and consequent under-editing have been mechanistically linked to motor-neuron death and TDP-43 pathology^11–20,32^.

**Figure 1.**
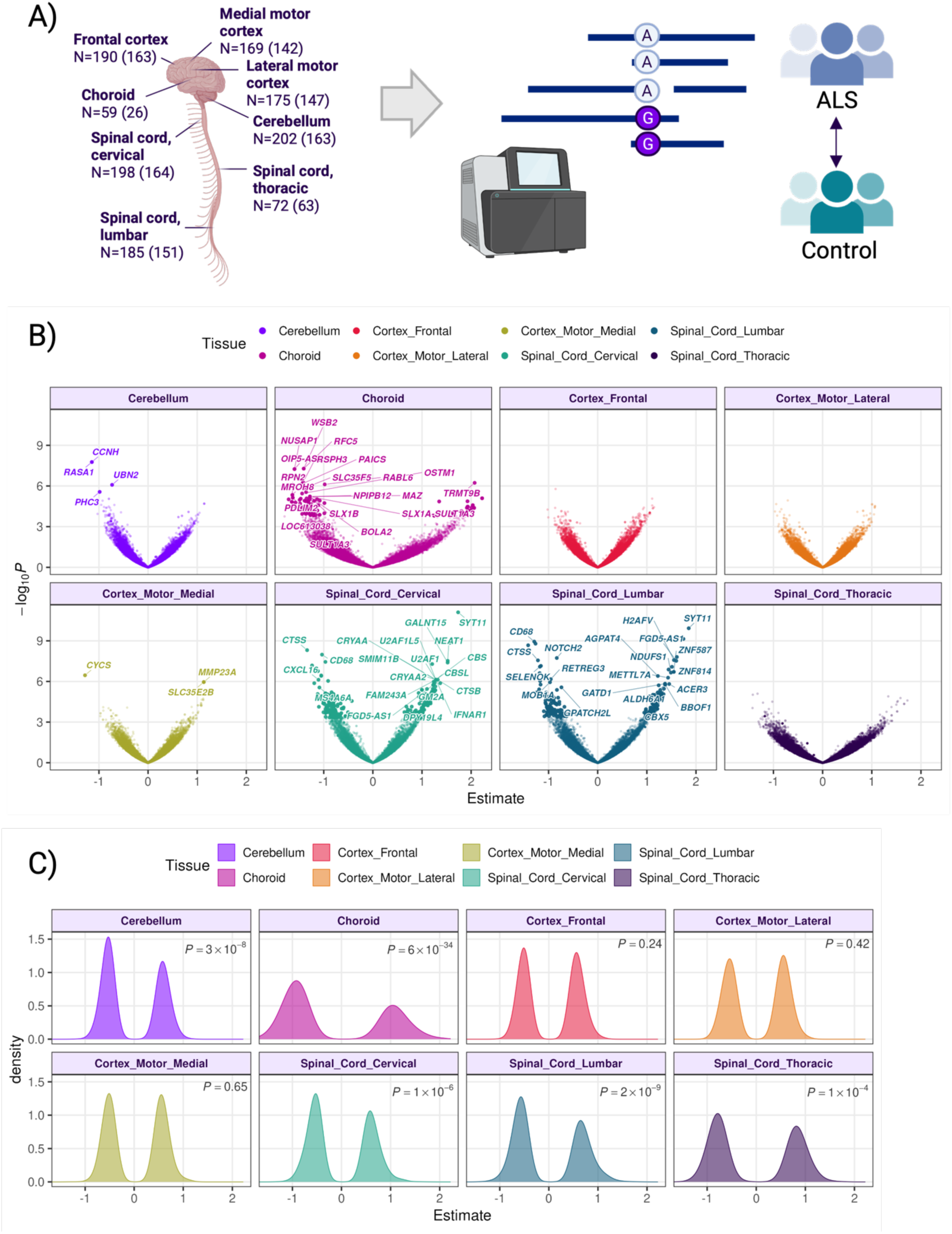
RNA editing is altered in ALS patient spinal cords. **(A)** Schematic of A-to-I editing detection from RNA-seq and the ALS vs. non-neurological control comparison. Case and control counts are shown for each tested tissue. Schematic elements were created with BioRender. **(B)** Volcano plots of differential editing for all 8 CNS regions. Representative significantly edited genes are labeled and larger points represent significant edQTL sites (FDR < 0.05) **(C)** Distribution of per-site differential-editing effect estimates by region, for each site with nominal *P*<0.05. Binomial test P-values are reported on the figures.

We then analyzed the differentially edited genes by Ingenuity Pathway Analysis. While no pathways reached significance after multiple testing correction, nominally significant pathways highlighted multiple disease-relevant themes (Supplementary File 2). The most reproducible signal was the cysteine-biosynthesis / transsulfuration pathway (*P*<0.05 six of eight tissues). This signal was driven by *CBS/CBSL*, the rate-limiting transsulfuration enzymes that detoxify homocysteine and generate hydrogen sulfide, in line with the elevated homocysteine reported in ALS patients^67^. Innate-immune and interferon programs were also recurrently represented (e.g. neutrophil degranulation, interferon-α/β and IL-10 signaling, PKR-mediated signaling), predominantly among spinal-cord and choroid genes, consistent with dsRNA sensing through PKR/EIF2AK2 driving the integrated stress response and neuroinflammation^68,69^. These observations align with the established role of dsRNA editing in restraining innate immune responses and suggest that impaired editing may contribute to ALS pathogenesis and progression^16–20^. Lastly, we detected editing changes in core RNA-splicing machinery (*U2AF1*, *SNRPD3*), echoing the significant role of splicing dysregulation and TDP-43-dependent cryptic-exon mis-splicing in ALS^70,71^. Together, these results nominate immune signaling, the integrated stress response and RNA processing as processes perturbed at the post-transcriptional level in ALS.

### Variants increasing RNA editing of DEGS1 3’ UTR are associated with ALS

Having shown that overall RNA editing is reduced in ALS, we asked whether variants influencing editing levels (edQTLs) are also associated with ALS risk. We mapped cis-edQTLs across Target ALS participants in eight tissues (N = 26–164, Fig. 2A). We tested 29,046–44,480 editing sites per tissue and identified significant edQTLs (FDR < 0.05) for up to 582 sites per tissue, with the most in cerebellum (582), cervical (567) and lumbar (424) spinal cord (Supplementary Table 2). To assess their relevance to disease, we performed genetic colocalization analysis between significant edQTL loci (FDR < 0.05) and ALS GWAS^3^ loci with suggestive association (*P* < 5x10^-6^). Two loci colocalized with posterior probability of a shared causal variant (PP.H4) > 0.80: *DEGS1* and *FNBP1* (Table 1, Supplemental Table 3). Neither loci reached genome-wide significance in the ALS GWAS (*P*>5x10^-8^), although *FNBP1* has previously linked to ALS through gene-based analyses^72^.

**Figure 2.**
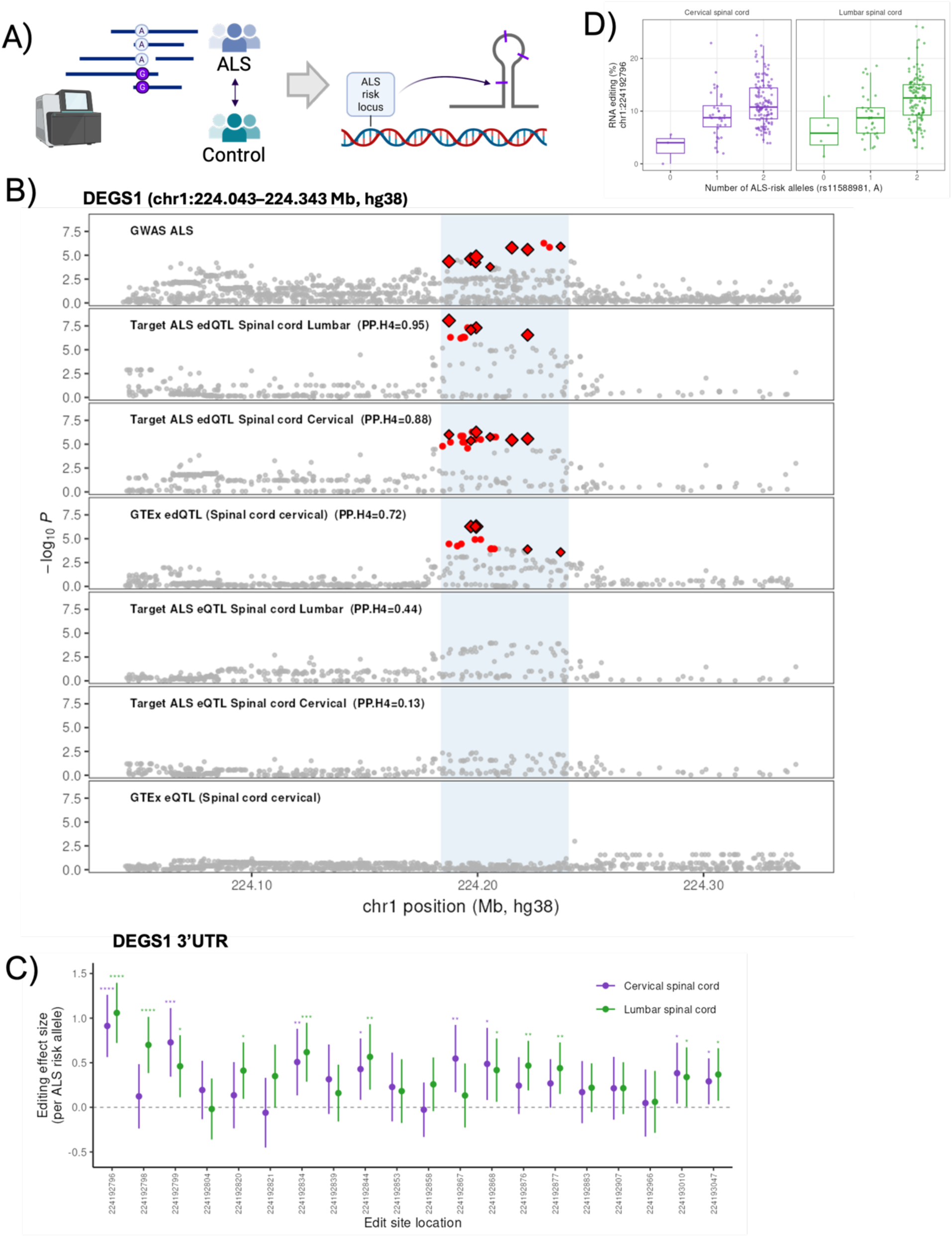
RNA editing of DEGS1 3’ UTR colocalizes with ALS GWAS risk variants. **(A)** Study design. A-to-I editing was quantified from RNA-seq from the Target ALS post-mortem cohort across eight CNS regions. Leveraging whole-genome sequencing (WGS), edQTL were identified and colocalized with GWAS risk loci. Numbers shown represent the total number of samples included with RNA-seq data, with total number included for edQTL mapping in parenthesis (matched WGS and RNA-seq). Schematic elements were created with BioRender. **(B)** Stacked locus plot for the *DEGS1* region (chr1:224.043–224.343 Mb) showing the ALS GWAS, Target ALS lumbar and cervical spinal cord and GTEx spinal cord. Fine-mapped credible-set variants are shown in red. Colocalization posterior probabilities (PP.H4) are reported on the figure. Diamonds mark variants colocalizing with the GWAS signal, with size related to the SuSiE-coloc variant posterior. **(C)** edQTL effect size, relative to the ALS-risk allele (rs11588981, A), at every *DEGS1* editing site (20 sites total). Error bars represent the 95% confidence interval. Asterisks denote nominal significance. **(D)** Fraction of RNA edited reads at the lead site versus the number of ALS-risk alleles (rs11588981, A) in cervical and lumbar spinal cord. Boxes show median and interquartile range. Samples with >10 counts at the sites are included

**Table 1.**
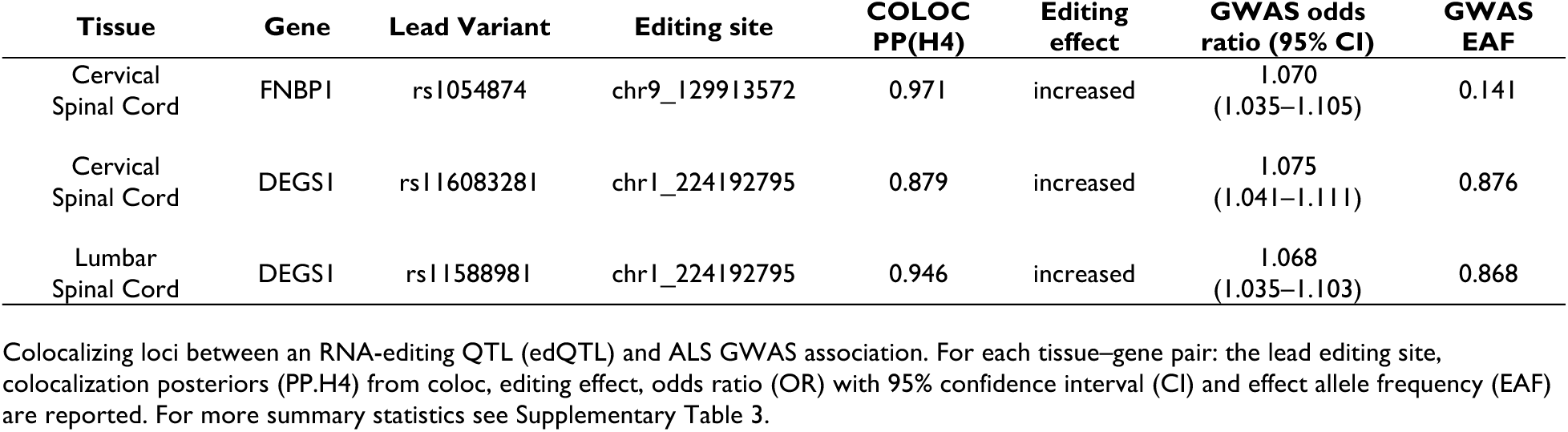
Top colocalizing edQTL with ALS GWAS.

*FNBP1* encodes an F-BAR domain protein that senses and generates membrane curvature to drive clathrin-mediated endocytosis and actin polymerization at synaptic membranes^73,74^, consistent with a role in motor neuron synaptic integrity within the spinal cord. Notably, a separate study using molecular QTL mapping performed directly in postmortem ALS spinal cord nominated *FNBP1* as a risk gene acting through splicing and expression changes^37^, providing tissue-matched evidence that parallels our own spinal cord colocalization findings. At *FNBP1*, editing colocalized with ALS risk in cervical spinal cord (PP.H4 = 0.97, Table 1, Supplemental Table 3, Supplementary Figure 2). Each ALS-risk allele of rs1054874 associated with +4.5% editing at the lead site, located in the intron of *FNBP1* (chr9:129,913,573). At *DEGS1*, the ALS-risk allele was associated with increased 3′ UTR editing and higher ALS risk in both lumbar (PP.H4 = 0.95) and cervical (PP.H4 = 0.88) spinal cord (Fig. 2B). On average, each copy of the lead edQTL (rs11588981) was associated with ∼3% higher editing at the lead 3′ UTR site (chr1:224,192,796; lumbar +3.1%, cervical +2.8% per allele; Fig. 2C-D, Supplementary Figure 1). Fine-mapping–based colocalization^62^ (SuSiE-coloc, methods) corroborated both loci (PP.H4 = 0.95 for *DEGS1* and 0.98 for *FNBP1*).

Given the uncharacterized association between ALS and DEGS1, a ceramide synthesizing enzyme, and growing evidence of elevated ceramides in ALS²³⁻³⁴, we investigated how *DEGS1* RNA editing affects ALS risk. We note, however, that three of the four most strongly ALS-associated SNPs at the *DEGS1* locus were absent from Target ALS WGS from the in-sample fine-mapping as they were removed during quality control, limiting our ability to identify shared causal variants. To determine whether the observed editing associations reflected underlying changes in gene expression, we mapped expression QTLs (eQTLs) in the same Target ALS tissues in which we identified colocalizing edQTLs. Unlike the robust colocalization observed between ALS risk and *DEGS1* RNA editing, *DEGS1* gene expression showed only weak colocalization with the risk locus, with a modest trend toward reduced gene expression in lumbar spinal cord (eQTL PP.H4 = 0.44 lumbar, 0.13 cervical). Similar findings were obtained using GTEx spinal cord eQTL data^63^ (Fig. 2B). Thus, the genetic association with *DEGS1* RNA editing appears to be independent of effects on *DEGS1* expression (Fig. 2B, 3A), highlighting RNA editing of *DEGS1* as the molecular trait associated with ALS risk.

### ALS-associated editing of the *DEGS1* 3’ UTR affects protein translation

DEGS1 catalyzes the conversion of dihydroceramide to ceramide, a key precursor for several bioactive sphingolipids^26,27^. To determine whether *DEGS1* gene expression is altered in ALS, we analyzed bulk RNA sequencing data from spinal cord tissue of sporadic and familial ALS cases from Target ALS and found no significant changes in *DEGS1* mRNA expression (Fig. 3A), complementing our findings of weak colocalization between eQTL and ALS risk variants (Fig. 2B). However, the absence of transcriptional changes does not exclude the possibility that DEGS1 protein abundance or activity may be altered through post-transcriptional regulatory mechanisms.

**Figure 3.**
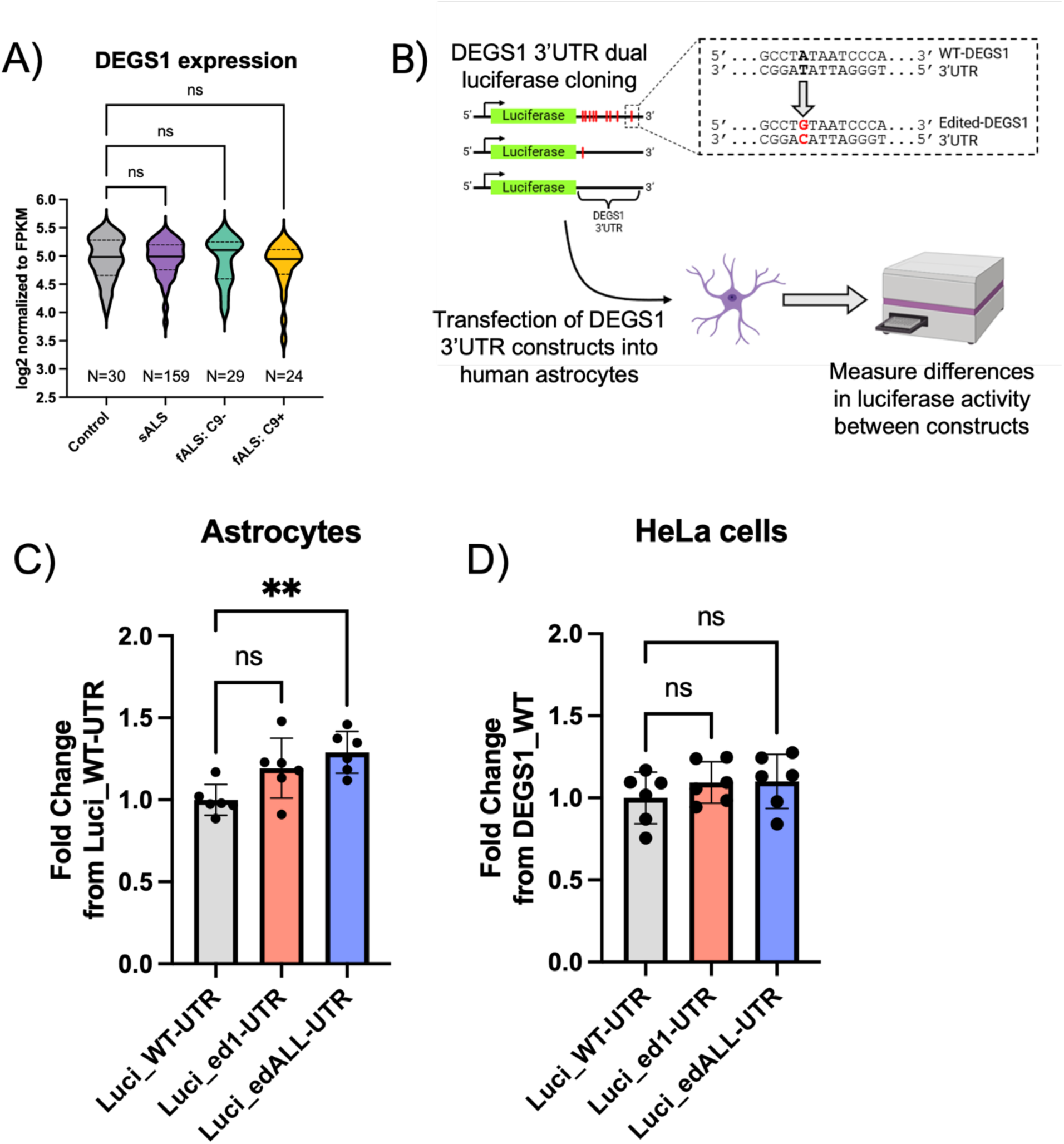
RNA editing of the DEGS1 3’UTR increases protein translation in human astrocytes. **(A)** *DEGS1* expression of control (N=30), sporadic (N=159) and familial ALS with (C9+; N=29) or without (C9-;N=24) *C9ORF72* mutation from Target ALS post-mortem spinal cords. **(B)** Experimental design. Luciferase constructs with either unedited or edited *DEGS1* 3’ UTR: wild type (Luci_WT-UTR), the most strongly affected edit site edited (Luci_ed1-UTR; chr1:224192796) and all identified edit sites edited (Luci_edALL-UTR). For each edit site A was switched to G. Constructs were transfected into human astrocytes and luciferase activity was measured. Schematic elements were created with BioRender. **(C and D)** Differences in luciferase activity across the different constructs in human astrocytes **(C)** or HeLa cells **(D)**. N=6 per group. ** denotes p < 0.01 using ordinary one-way ANOVA with Dunnett’s multiple comparisons test.

Our analysis identified increased RNA editing at specific sites within the *DEGS1* 3′ UTR that were linked to elevated ALS risk (Fig. 2C, Supplementary Figure 1). Given the well-established role of 3’ UTRs in regulating protein expression, we hypothesized that these editing events may functionally influence DEGS1 protein expression. To test this possibility, we evaluated the translational effects of the edited and unedited 3′ UTR constructs in astrocytes. Astrocytes were selected due to their critical roles in lipid metabolism and signaling, and neuroinflammatory processes, including the production and propagation of pro-inflammatory cytokines, all of which are implicated in ALS pathology^75–78^. Moreover, ceramide synthase inhibition resulted in a greater reduction in *de novo* ceramide synthesis in glial cells than in motor neurons, further justifying the use of astrocytes as the primary cellular model in this study^23^. We used a dual luciferase assay to determine whether the presence of edited *DEGS1* 3’ UTR could result in changes in luciferase activity as a measure of protein expression. Three different versions of the unedited or A-to-G edited 3’ UTR of *DEGS1* were cloned into a dual luciferase vector: (1) Wildtype (Luci_WT-UTR), (2) 3’ UTR with the most affected site edited (Luci_ed1-UTR, chr1_224192796), (3) 3’ UTR with all sites edited (Luci_edALL-UTR) (Fig. 3B, Supplementary File 3). Transfection of Luci_edALL-UTR into human astrocytes resulted in a significant increase in luciferase activity compared to Luci_WT-UTR while transfection of Luci_ed1 showed no effect (Fig. 3C). In contrast, transfection of these constructs into HeLa cells showed no difference in luciferase activity relative to Luci_WT-UTR (Fig. 3D). Together, this shows that RNA editing at the 3’ UTR of *DEGS1* may increase expression of its upstream protein as exhibited by the increase in luciferase activity and that this regulatory effect might be context or cell type specific.

### Knockout of DEGS1 perturbs synthesis of ceramides and downstream sphingolipids

To elucidate the extent of DEGS1 activity on sphingolipid metabolism, we used CRISPR/Cas9 to generate DEGS1 KO iPSCs and used lipidomic profiling to determine changes in lipid levels (Fig. 4A). Individual clones were isolated and propagated followed by western blotting for DEGS1 to confirm successful knockout (Fig. 4B). Several of the clones showed little to no DEGS1 expression compared to control transfected or WT HEK293T cells. For subsequent experiments, clone D6 was arbitrarily selected for further analysis (Fig. 4B).

**Figure 4.**
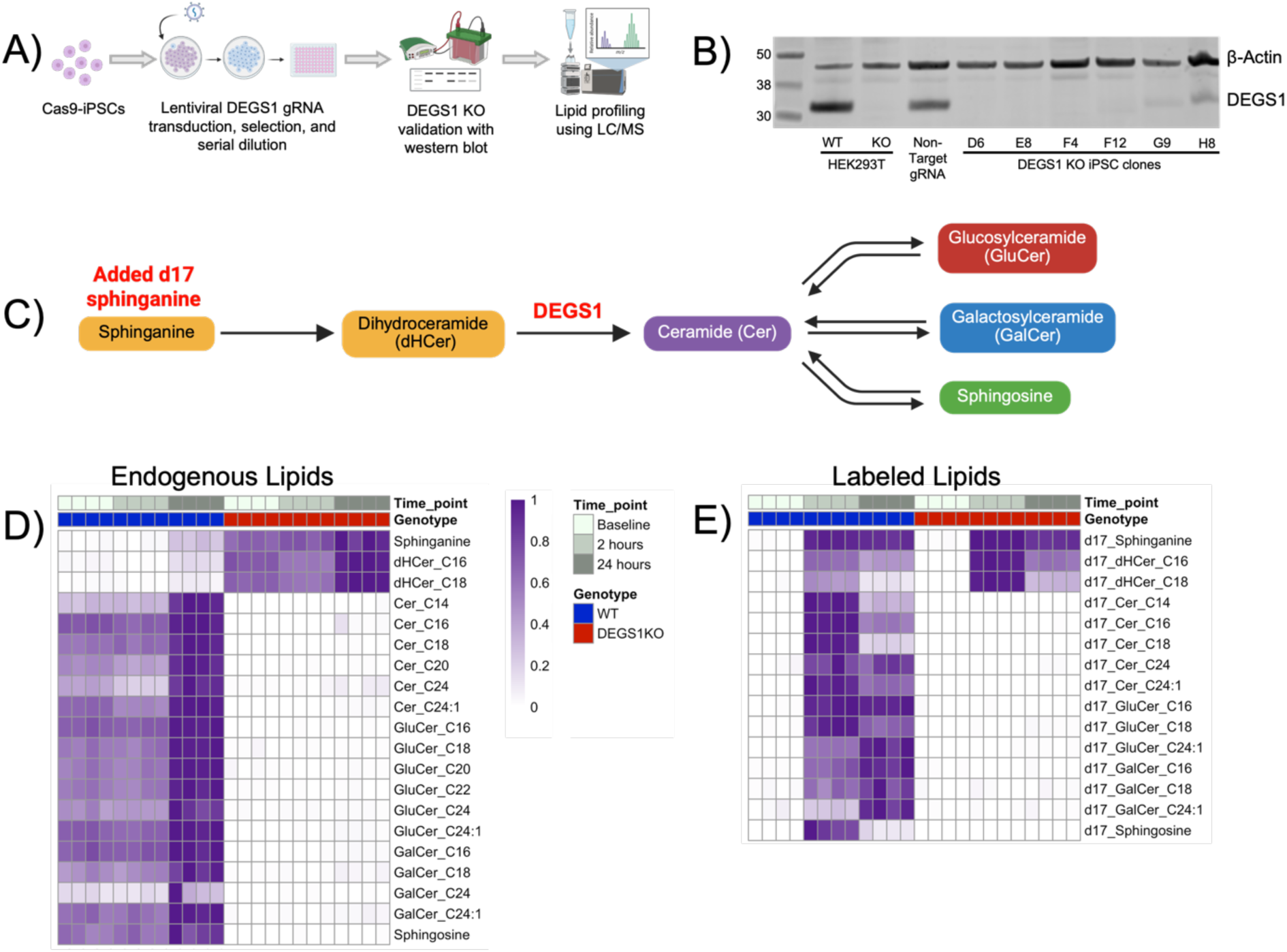
DEGS1 is necessary for sphingolipid metabolism. **(A)** Schematic of generation and validation of DEGS1 KO iPSCs. Schematic elements were created with BioRender. **(B)** Western blot of showing several clones with decrease or loss of DEGS1 protein compared to non-targeting gRNA controls. Clone D6 was chosen for lipid profiling and used for subsequent experiments. **(C)** schematic of sphingolipid pathway analyzed. d17-sphinganine was used for lipid labeling experiments. **(D and E)** d17-sphinganine was added to cells for 2 or 24 hours in WT (blue) or DEGS1 KO iPSCs (red). Baseline had no d17-sphinganine added. Cells were used for lipidomic analysis of both endogenous lipids **(D)** and labeled lipids containing d17 sphingoid base **(E)**. For each lipid analyzed values were scaled with the highest value equal to 1 (dark purple) and lowest to 0 (white).

To evaluate changes in *de novo* sphingolipid synthesis, we added the ceramide precursor lipid, sphinganine, with an artificial 17-carbon chain length (d17-sphinganine). To capture changes in sphingolipids over time, d17-sphinganine was added to WT or DEGS1 KO iPSCs for either 2 or 24 hours and LC/MS was used to profile the following lipids of various carbon chain lengths: sphinganine, dihydroceramide (dHCer), ceramide (Cer), glucosylceramide (GluCer), galactosylceramide (GalCer) and sphingosine. These lipids were selected due to their proximity to DEGS1 activity and their dysregulation in the context of sporadic ALS^30–32,79^ (Fig. 4C). Consistent with its role in sphingolipid metabolism, endogenous Cer, GluCer, GalCer and sphingosine synthesis was abolished across all time points in DEGS1 KO iPSCs compared to WT; furthermore, the precursor lipids sphinganine and dHCer accumulated in DEGS1 KO cells (Fig. 4D). These trends were observed in labeled sphingolipids which incorporated the d17-backbone: d17-Cer, d17-GluCer, d17-GalCer and d17-sphingosine synthesis was absent across all time points and d17-sphinganine and d17-dHCer were increased in DEGS1 KO iPSCs compared to WT (Fig. 4E). These findings underscore the instructive role of DEGS1 in orchestrating the sphingolipid environment, suggesting that changes in DEGS1 expression may directly alter lipid levels.

### ALS-associated editing of the 3’UTR increases DEGS1 expression and modifies ceramide levels

We demonstrated that ALS-associated edits at the 3’ UTR of *DEGS1* leads to increased luciferase activity (Fig. 3C), signifying that these edits may increase the expression of its upstream protein in astrocytes. To test this, we generated iPSC-derived DEGS1 KO astrocytes and transfected them with human DEGS1 containing either edited or unedited 3’ UTRs (Fig. 5A). Successful differentiation of iPSCs into astrocytes was confirmed through staining with the astrocyte markers S100β and CD44^80,81^ (Supplementary Figure 3A). These astrocytes also preserved the ceramide synthesis deficits previously seen in iPSCs (Supplementary Figure 3B). DEGS1 KO astrocytes were transfected with the following constructs: human *DEGS1* containing 3’ UTR with no edits (DEGS1_WT-UTR), most affected site edited (DEGS1_ed1-UTR), or all sites edited (DEGS1_edALL-UTR). These constructs were co-transfected with GFP to account for differences in transfection efficiency. Transfection of DEGS1_edALL-UTR into DEGS1 KO astrocytes resulted in a significant increase in DEGS1 protein expression relative to DEGS1_WT-UTR and DEGS1_ed1-UTR (Fig. 5B). As DEGS1_WT-UTR and DEGS1_ed1-UTR showed minimal differences and that editing frequency is variable across all sites (Fig. 2C, Fig. 3C, Fig.5B, Supplementary Figure 1), subsequent analyses focused on comparing the fully edited 3′ UTR with the wild-type 3′ UTR.

**Figure 5.**
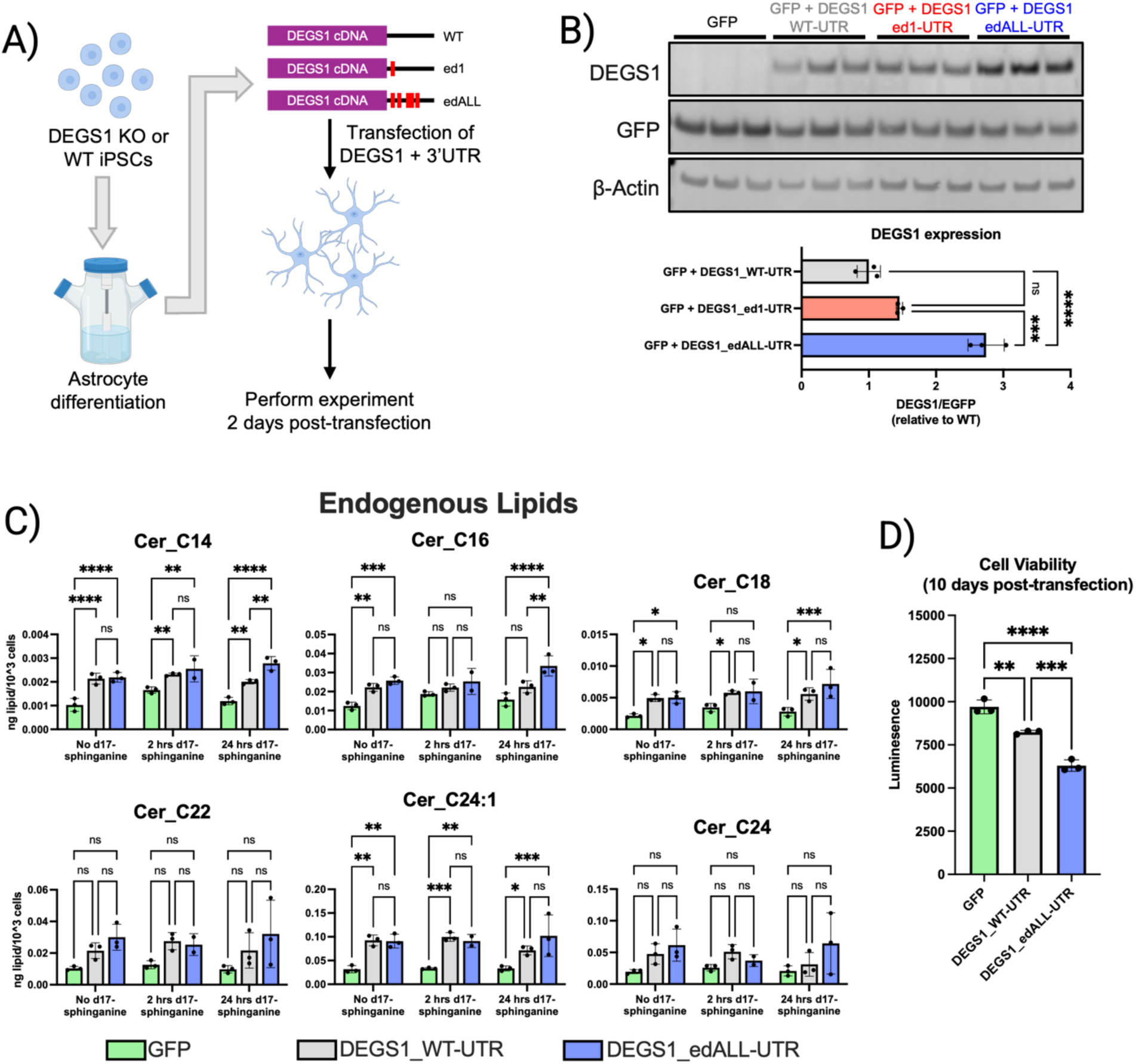
Edited DEGS1 3’ UTR increases protein translation and promotes ceramide synthesis leading to decreased cell viability in iPSC-Astrocytes. **(A)** Non-target gRNA or DEGS1 KO iPSCs were differentiated into astrocytes and transfected with human DEGS1 with: (1) DEGS1_WT-UTR; (2) the most affected edit site, DEGS1_ed1-UTR; (3) all identified edit sites, DEGS1_edALL-UTR. Experiments were performed 2 days after transfection. Schematic elements were created with BioRender. **(B)** DEGS1 protein expression when DEGS1 transcripts with WT or unedited 3’ UTR was transfected. DEGS1 values were normalized to their corresponding EGFP values. All groups were compared to DEGS1_WT-UTR, N=3 per group. **(C)** Targeted lipid profiling of DEGS1 KO astrocytes following transfection with GFP, DEGS1_WT-UTR, or DEGS1_edALL-UTR for endogenous ceramides at 2 hours and 24 hours after d17-sphinganine addition 2 days post-transfection. d17-sphinganine was added to both groups at the same time and collected at different times. The levels of C14, C16, C18, C22, C24:1, or C24 acyl chain length ceramides were normalized to total cell count. N=2 for 2 hr d17-sphinganine, edALL group; N=3 for all other groups. **(D)** Cell viability of DEGS1 KO astrocytes transfected with GFP, DEGS1_WT-UTR or DEGS1_edALL-UTR after 10 days. * denotes p < 0.05, ** denotes p < 0.01, *** denotes p < 0.001, **** denotes p < 0.0001 using ordinary one-way ANOVA with Dunnett’s multiple comparisons test (B, D) or two-way ANOVA with Tukey’s multiple comparisons test.

To determine the impact of RNA editing of the 3’ UTR of *DEGS1* on ceramide levels we performed targeted lipid profiling of endogenous and d17-labeled ceramides of different acyl chain lengths (C14, C16, C18, C22, C24:1, and C24) in DEGS1 KO astrocytes expressing DEGS1_WT-UTR or DEGS1_edALL-UTR. Expression of either DEGS1_WT-UTR or DEGS1_edALL-UTR in DEGS1 KO astrocytes resulted in a significant increase in several endogenous and d17-labeled ceramide species compared to GFP transfected controls (Fig. 5C). Importantly, expression of DEGS1_edALL-UTR resulted in significantly elevated endogenous C14 and C16 ceramide levels 24 hours after d17-sphinganine stimulation relative to DEGS1_WT-UTR (Fig. 5C). No significant difference in d17-labeled ceramide levels were found between DEGS1_edALL-UTR and DEGS1_WT-UTR except for d17_Cer_C14 (Supplementary Figure 4). Given the established role of ceramide synthase 5/6-derived ceramides, Cer_C14 and Cer_C16, in promoting multiple cell death pathways^23,82–84^, we next examined whether expression of the edited 3′ UTR affected cell viability. Indeed, transfection of DEGS1_edALL-UTR lead to a significant decrease in cell viability relative to DEGS1_WT-UTR (Fig. 5D). How these edits increase DEGS1 protein translation remains unclear. *In silico* analysis shows the edit sites lie mainly within an RNA hairpin in the predicted *DEGS1* 3’ UTR secondary structure, suggesting they may alter secondary structure to affect protein expression, or modify miRNA binding sites, altering regulatory control of protein expression (Supplementary Figure 5, 6). See the Discussion for further consideration of these mechanisms. Together, these findings demonstrate that ALS-associated post-transcriptional editing of the *DEGS1* 3′ UTR can increase DEGS1 protein expression, alter ceramide metabolism and promote cell death.

## Discussion

Dysregulation of RNA editing has emerged as a potential mechanism in ALS, linking altered post-transcriptional regulation to motor neuron degeneration^11–20,32^. Here, we observed colocalization of ALS GWAS signals among common genetic variants associated with RNA editing, leading to increased RNA editing within the 3’UTR of *DEGS1*. We demonstrate that DEGS1 functions as a major regulator of multiple sphingolipid species in astrocytes and show that increased RNA editing within its 3′ UTR enhances protein expression, thereby increasing ceramide synthesis to promote cell death. Importantly, we show these effects occur in astrocytes, a cell type increasingly recognized as a driver of ALS progression through maladaptive metabolic and inflammatory signaling, including dysregulated sphingolipid metabolism. Collectively, these results suggest that RNA editing of *DEGS1*, supported by enrichment of ALS GWAS signals, represents a previously unrecognized regulatory mechanism linking genetic variation to altered sphingolipid metabolism and ALS risk.

### Differential effects on endogenous and d17-labeled ceramides

We observed a significant increase in endogenous Cer_C14 and Cer_C16 after 24 hours post-d17 stimulation, but not their corresponding d17-labeled species, following re-expression of *DEGS1* containing the edited 3′ UTR compared with the WT 3′ UTR in DEGS1 KO iPSC-derived astrocytes. Endogenous ceramides represent the cumulative balance of de novo synthesis, salvage pathways, and conversion into downstream sphingolipids during the several days following DEGS1 re-expression. In contrast, d17-labeled species are derived exclusively from the exogenous d17-sphinganine precursor during the 2-hour or 24-hour labeling periods. This distinction suggests that restoration of DEGS1 allows ceramide levels to re-equilibrate to a new steady state over time, whereas the labeled lipid pool captures only those lipids synthesized from the d17-backbone during the labeling window immediately preceding sample collection. Furthermore, because d17-sphinganine was supplied as a single pulse rather than by continuous infusion, the resulting d17-labeled lipid species primarily reflect the fate and distribution of newly synthesized lipids over time and are not well suited to direct comparisons of steady-state metabolic flux between the WT and edited 3′ UTR conditions.

The absence of differences in d17-labeled ceramides despite differences in endogenous ceramide levels may also reflect compensatory adaptations within the sphingolipid metabolic network. Re-expression of DEGS1 perturbs a tightly regulated pathway, and the 2 to 3 days between transfection and sample collection provide sufficient time for homeostatic responses to occur. Such adaptations could involve serine palmitoyl transferase and its ORMDL regulators, which control entry into *de novo* sphingolipid synthesis; ceramide synthase isoforms, which determine acyl-chain composition; and ceramidases, sphingomyelin synthases, and glucosylceramide synthase, which regulate ceramide turnover and conversion into downstream sphingolipids. Furthermore, enzymes acting at multiple points within the sphingolipid pathway, including ACER3, SGMS2, and GM2A, were also associated with differential RNA editing, highlighting the possibility of these edit sites modifying protein expression of their respective genes (Supplementary File 1). The presence of multiple independent editing sites within ACER3, which directly regulates ceramide catabolism, is particularly noteworthy and suggests that RNA editing may influence sphingolipid homeostasis beyond DEGS1. Changes at any of these regulatory nodes could alter steady-state ceramide abundance or redistribute sphingolipid flux throughout the pathway. Because DEGS1 protein levels differ between cells expressing the edited and unedited *DEGS1* 3′ UTRs, the magnitude of these compensatory responses may likewise differ between conditions. In addition, our lipidomic analysis was restricted to ceramide species and therefore cannot exclude the possibility that alterations in sphingolipid metabolism were manifested in downstream sphingomyelin, sphingosine, or hexosylceramide pools rather than in ceramide itself.

An additional limitation is that all measurements were performed in transiently transfected cultures. Consequently, the data represent population averages across a heterogeneous mixture of untransfected DEGS1-null cells and transfected cells expressing varying levels of DEGS1. Untransfected cells are expected to dilute construct-dependent effects toward the null phenotype, while differences in transfection efficiency introduce experimental variability unrelated to the underlying biology. Together, these factors may reduce the sensitivity of comparisons between the edited and unedited *DEGS1* 3′ UTRs.

### Mechanisms of translational regulation

The exact mechanism on how RNA editing at these specific sites on the 3’ UTR of *DEGS1* and how it modulates protein expression is still unresolved. One explanation could be that RNA editing of the 3’ UTR could disrupt its secondary structure thus potentially disrupting inhibitory stem-loop structures that are ultimately targeted by RNA binding protein to repress translation (Supplementary Figure 5). Indeed, this has been demonstrated in mesothelioma wherein RNA editing at the 3’ UTR of *RBM8A* altered its secondary structure and elevated RBM8A protein levels by preventing RNA binding protein-dependent translational repression^85^. Another potential mechanism could be changes in miRNA binding sites. It was found that RNA editing of the 3’ UTR of *ARHGAP26* disrupted miRNA binding sites which lead to upregulation of the protein and subsequent modulation of its downstream partner, *RhoA*^86^. As an exploratory analysis, we performed a similar analysis to determine if miRNA binding sites at the 3’ UTR were altered by RNA editing.

We analyzed three *DEGS1* 3′ UTR variants (WT, ed1, edALL) for predicted binding of mature human miRNAs (miRBase). For each UTR, miRNA seed matches were identified using 8mer, 7mer-m8 and 7mer-A1 criteria (Supplementary Figure 6A). Edited 3’ UTRs were compared with WT to identify miRNA binding sites that were lost or gained. The single edit in ed1 resulted in the loss of one miRNA binding site, whereas edALL, which contains multiple edits, led to the loss of 21–41 sites and the gain of 4–9 sites, depending on seed-match stringency (Supplementary Figure 6B). We next determined ALS relevance by cross-referencing the Human microRNA Disease Database (HMDD v4.0)^66^ and PubMed for studies reporting each miRNA and its connection to ALS or motor neuron disease, narrowing the list to 13 miRNAs (Supplementary File 4). Mapping these miRNAs to the WT and edALL 3’ UTRs showed that several binding sites were lost upon RNA editing and sites that were retained exhibited reduced binding under more stringent seed-matching criteria (Supplementary Figure 6C, D). It is unknown whether the creation of new miRNA binding would directly modulate DEGS1 protein expression or act as a decoy for its natural targets. Nevertheless, this reveals that ceramide dysregulation in ALS can potentially be caused by aberrant miRNA activity via RNA editing.

### Variants associated with ALS risk and DEGS1 edQTLs

Suggestive ALS risk variants at the *DEGS1* locus colocalized with RNA editing of the gene. While the association was suggestive, DEGS1 resides in the sphingolipid pathway that is strongly linked to ALS. One variant resides within an intron of *DEGS1* (rs11588981). Intronic and untranslated regions commonly participate in the formation of double-stranded RNA structures that serve as ADAR substrates and modest sequence variation influences editing efficiency at nearby sites^87,88^. Moreover, RNA editing often occurs co-transcriptionally and is coupled to pre-mRNA processing, suggesting that intronic variation may affect editing through changes in RNA secondary structure or splicing^89^. The remaining SNPs are in intergenic regions (rs116083281, rs6426060) and within an intron non-coding gene (rs10916515), indicating that distal or indirect regulatory mechanisms may also contribute. Intergenic variants can tag enhancers or other regulatory elements that modulate transcriptional kinetics or chromatin state, factors that have been associated with variability in RNA editing levels^11^. In addition, non-coding RNAs are known to interact with ADAR enzymes and may influence editing by competing for ADAR binding or altering RNA–RNA interactions in cis or trans^88,90^. The associated SNPs are in strong linkage disequilibrium and 3 of the lead GWAS variants were not included in the Target ALS analysis as they failed quality control filters. Thus, statistical fine mapping of putative causal variants remains ambiguous. Nonetheless, these findings support a model in which RNA editing of *DEGS1* is influenced by its broader regulatory context.

### Cell type specificity of RNA editing

Our results demonstrate that RNA editing of the *DEGS1* 3′ UTR increases DEGS1 protein expression without altering mRNA levels, suggesting a post-transcriptional mechanism. This interpretation is supported by the lack of colocalization between DEGS1 expression quantitative trait loci (eQTLs) and ALS GWAS variants in the spinal cord, as well as by the absence of differential *DEGS1* gene expression between familial or sporadic ALS cases and healthy controls (Fig. 2B, Fig. 3A). While sphingolipid dysregulation is implicated in ALS, changes in *DEGS1* gene (Fig. 3A) or protein expression have not previously been reported. Prior studies of GluA2 Q/R site editing in sporadic ALS show cell type–specific differences, with altered editing in spinal motor neurons but not in Purkinje cells or motor cortex^12,13,15^, highlighting the cell-selective nature of RNA editing. We observe allele-dependent effects of genomic variants on individual *DEGS1* edit sites, with the most affected site showing 5–20% editing and others <5% (Fig. 2D, Supplementary Figure 1). These values may be underrepresented in bulk RNA-seq and may be enriched at the single-cell level, potentially revealing cell type-specific increases in DEGS1 protein expression.

We demonstrate that the fully edited 3′ UTR of *DEGS1* increases luciferase activity in astrocytes, but not in HeLa cells (Fig. 3D). Together, these findings suggest that dysregulated sphingolipid metabolism in astrocytes may contribute to ALS progression, a concept supported by several prior studies. Astrocytes are central regulators of lipid storage, processing, and signaling. Disruption of these functions can promote oxidative stress, mitochondrial dysfunction, activation of neuronal cell death pathways and inflammatory signaling all of which are phenotypes in ALS^77,78,91^. Notably, work in Drosophila has shown that DEGS1 facilitates the formation of intraluminal vesicles and the release of exosomes into the extracellular space^92^. This raises the possibility that astrocytes may propagate toxic factors, contributing to the spread of ALS pathology across cells and regions. However, whether sphingolipid dysregulation in ALS is primarily driven by astrocytes remains unclear. Further investigation of sphingolipid metabolism in other cell types, including motor neurons and microglia, will be essential for a more comprehensive understanding of sphingolipid dysregulation in ALS progression.

## Data availability

All data generated or analyzed that support the findings of this study are available from the corresponding author upon reasonable request.

## Competing interests

Alan Guo, Samuel Lessard, Lilu Guo, Melody Li, Disha Sood, Rachel Passaro, Jeremy Huang, Clement Chatelain, Emanuele de Rinaldis, Shameer Khader, Bailin Zhang, James C. Dodge and Steven Rodriguez are employees of Sanofi and may hold stock/options in the company.

## Supplementary Tables

**Supplementary Table 1.**
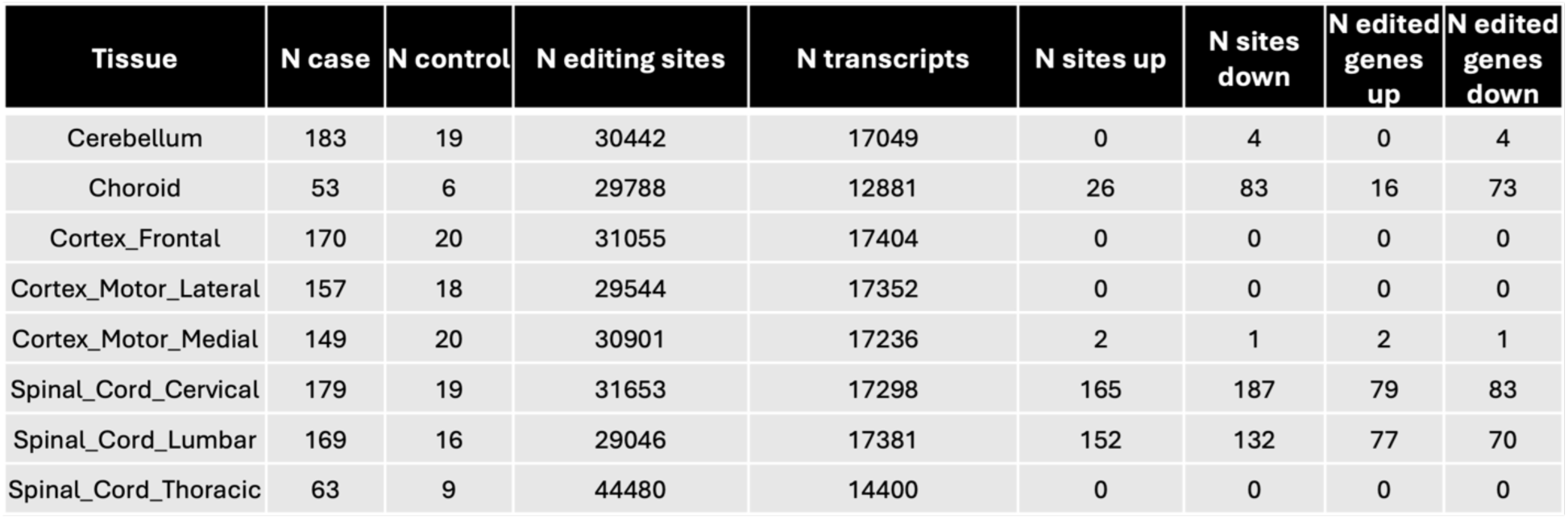
RNA-editing quantification and differential editing by tissue. Numbers of ALS cases and controls and number of quantified editing sites and transcripts per CNS region. Number of differentially edited sites are shown (ALS vs. control, FDR < 0.05), separated by directionality, as well as the corresponding number of genes. Positions are GRCh38-based.

**Supplementary Table 2.** Number of significant cis-edQTLs separated by tissue. Table reports the number of samples used to quantify edQTL (samples with matched RNAseq and WGS data), total number of sites tested and number of significant edQTL (permutation FDR < 0.05).

| Tissue | N samples | N sites tested | N sig edQTL (FDR < 0.05) |
| --- | --- | --- | --- |
| Cerebellum | 163 | 30442 | 582 |
| Choroid | 26 | 29788 | 0 |
| Cortex_Frontal | 162 | 31055 | 508 |
| Cortex_Motor_Lateral | 147 | 29544 | 405 |
| Cortex_Motor_Medial | 142 | 30901 | 373 |
| Spinal_Cord_Cervical | 164 | 31653 | 567 |
| Spinal_Cord_Lumbar | 151 | 29046 | 424 |
| Spinal_Cord_Thoracic | 65 | 44480 | 147 |

**Supplementary Table 3.** Top colocalizing edQTL with ALS GWAS (expanded) Expanded table from Table 1. Colocalizing loci between an RNA-editing QTL (edQTL) and ALS GWAS association. For each tissue–gene pair: the lead editing site, colocalization posteriors (PP.H4) from coloc and SuSiE-coloc are reported, as well as the summary statistics for the lead edQTL (smallest P-value at the locus) and summary statistics of the same variant in the GWAS. Directionality is concordant when the direction of effect of the edQTL and GWAS variants are in the same direction. Positions are GRCh38-based. SE: Standard error; OR: Odds ratio; CI: Confidence interval; EAF: Effect allele frequency.

| Tissue | Gene | Editing site | COLOC PP(H4) | SuSiE PP(H4) | Lead edQTL variant | Other allele | Effect allele | edQTL FastQTL q-value | edQTL beta | edQTL SE | edQTL P | GWAS OR | GWAS 95% CI | GWAS P | GWAS EAF | Directionality |
| --- | --- | --- | --- | --- | --- | --- | --- | --- | --- | --- | --- | --- | --- | --- | --- | --- |
| Spinal_Cord Cervical | FNBP1 | chr9_129913572 | 0.971 | 0.978 | rs1054874 | T | C | 0.00302 | 0.864 | 0.145 | 1.79E-08 | 1.07 | 1.035–1.105 | 4.54E-05 | 0.141 | Concordant |
| Spinal_Cord Lumbar | DEGS1 | chr1_224192795 | 0.946 | 0.954 | rs11588981 | G | A | 0.00161 | 1.059 | 0.172 | 8.66E-09 | 1.068 | 1.035–1.103 | 4.39E-05 | 0.868 | Concordant |
| Spinal_Cord Cervical | DEGS1 | chr1_224192795 | 0.879 | 0.953 | rs116083281 | A | G | 0.0304 | 0.955 | 0.182 | 5.28E-07 | 1.075 | 1.041–1.111 | 1.41E-05 | 0.876 | Concordant |

## Supplementary Figures

**Supplementary Figure 1:**
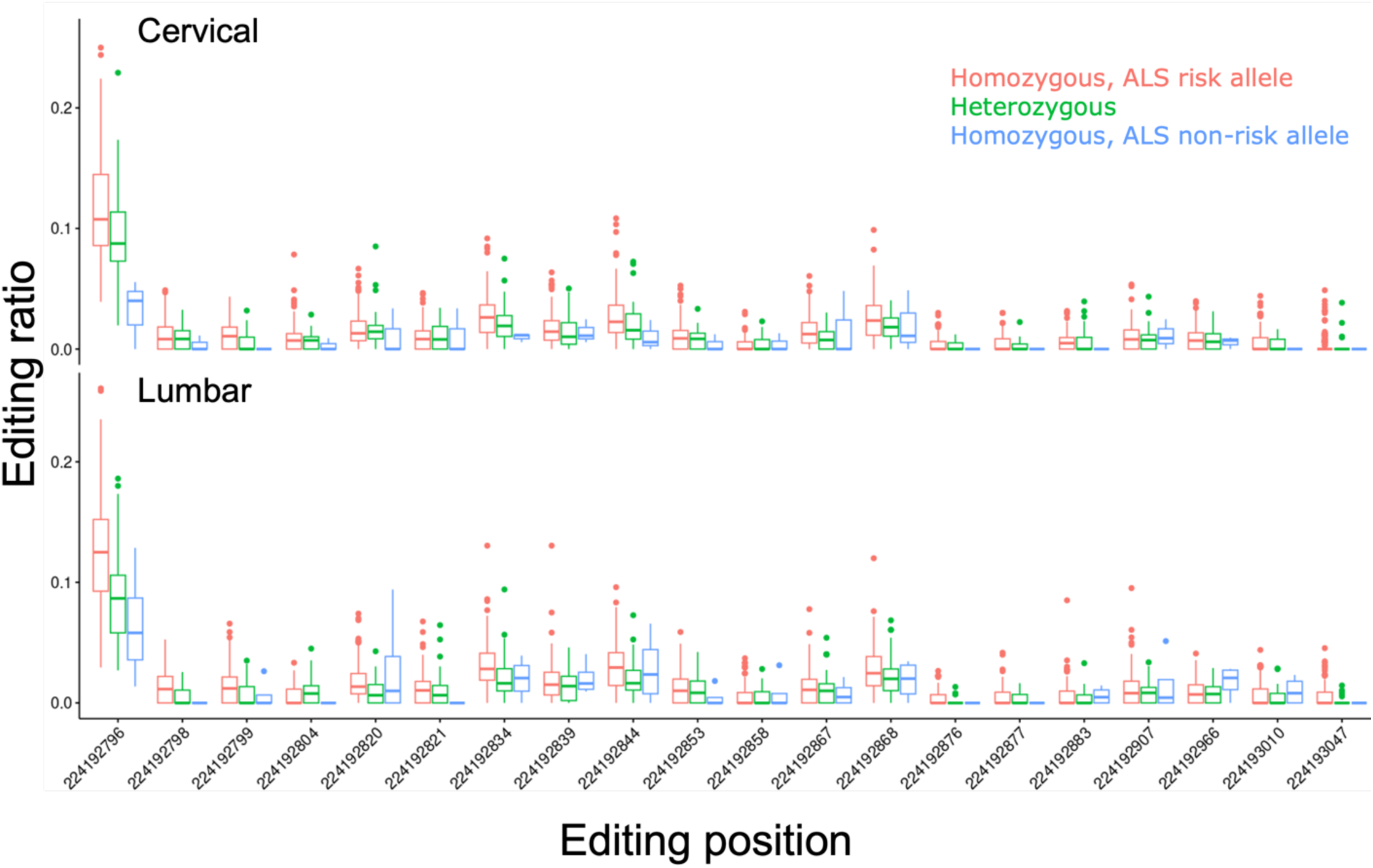
ALS risk allele affects DEGS1 RNA editing frequency. Fraction of RNA edited reads at all 20 *DEGS1* 3’UTR site versus the number of ALS-risk alleles (rs11588981, A) in cervical and lumbar spinal cord. Boxes show median and interquartile range. Samples with >10 counts at the sites are included.

**Supplementary Figure 2.**
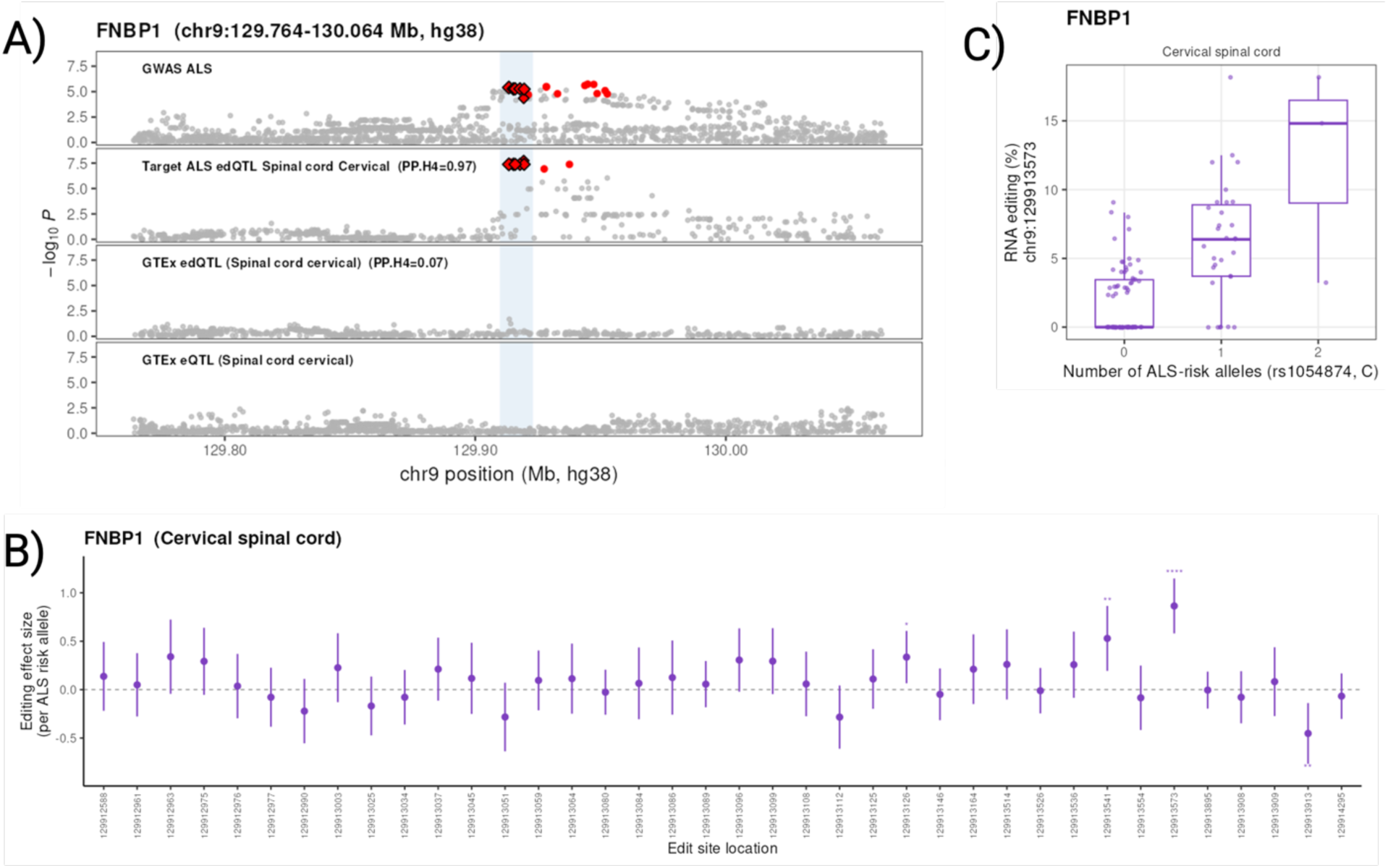
Colocalization of *FNBP1* editing-QTL with ALS risk. **(A)** Stacked locus plot for the *FNBP1* region (chr9:129.764–130.064 Mb) for the ALS GWAS, Target ALS edQTL and eQTL in cervical spinal cord and corresponding GTEx edQTL and eQTL (cervical). Fine-mapped credible-set variants are shown in red. Colocalization posterior probabilities (PP.H4) are reported on the figure. Diamonds mark variants colocalizing with the GWAS signal, with size related to the SuSiE-coloc variant posterior. **(B)** edQTL effect size, relative to the ALS-risk allele (rs1054874, C), at *FNBP1* gene-body editing sites around the lead site (204 sites, cervical spinal cord). Error bars represent the 95% confidence interval. Asterisks denote nominal significance. **(C)** Fraction of RNA edited reads at the lead site versus the number of ALS-risk alleles in cervical spinal cord. Boxes show median and interquartile range. Samples with >10 counts at the sites are included.

**Supplementary Figure 3.**
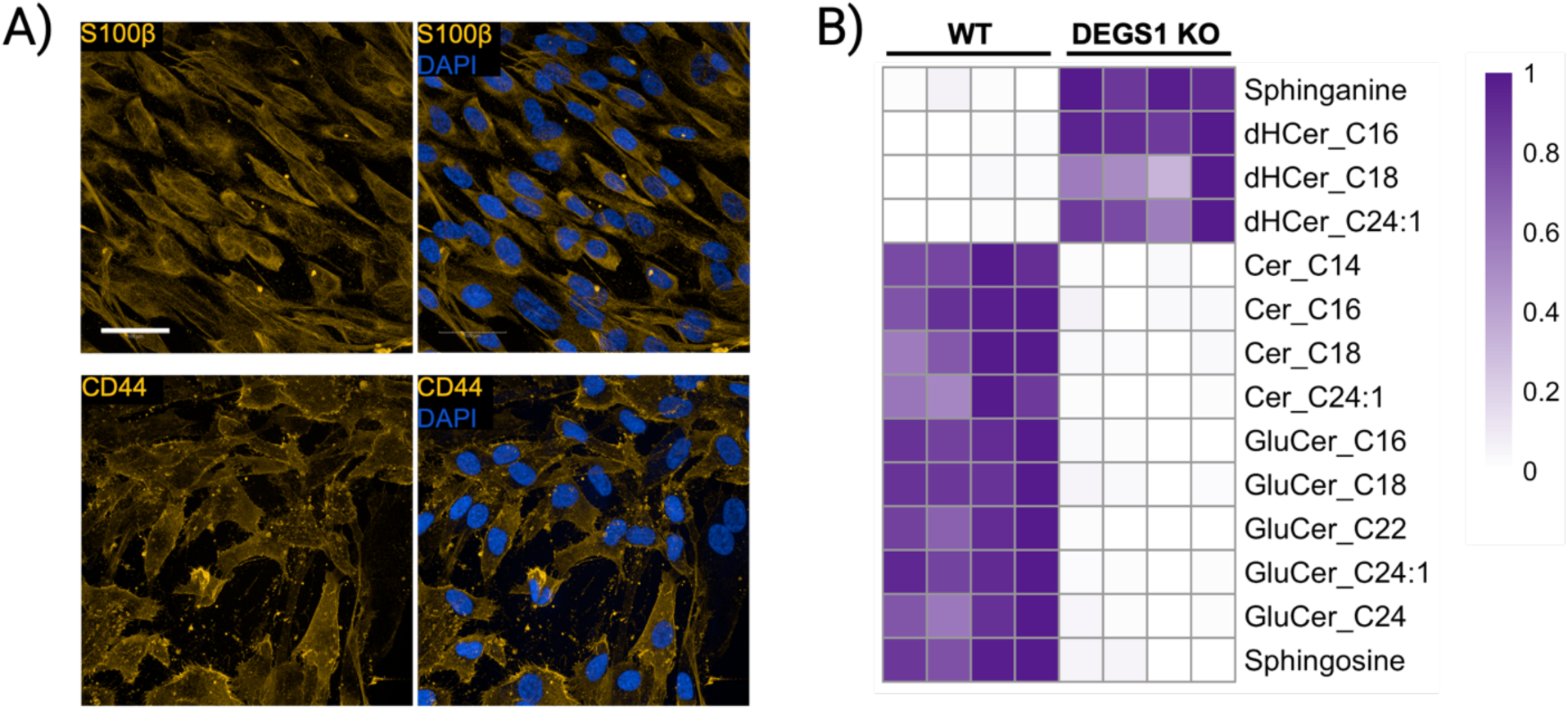
DEGS1 KO iPSC-derived astrocytes maintain sphingolipid loss. **(A)** Representative images showing iPSC-differentiated astrocytes were positive for astrocyte markers, S100β and CD44. **(B)** DEGS1 KO astrocytes show disrupted sphingolipid synthesis compared to WT.

**Supplementary Figure 4.**
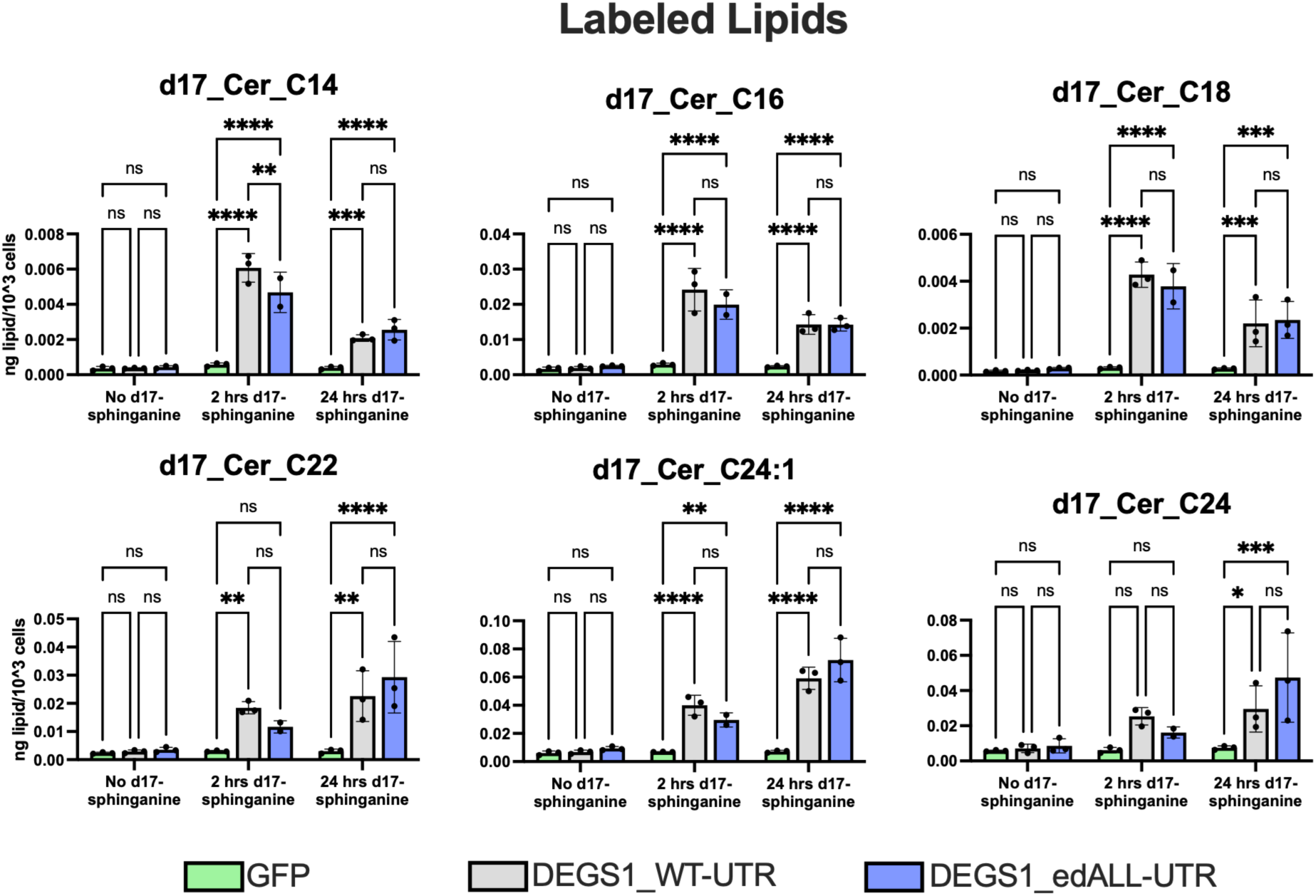
Edited DEGS1 3’ UTR modulates d17-labeled ceramide synthesis. Same samples as in Fig 5C. Targeted lipid profiling of DEGS1 KO astrocytes following transfection with GFP, DEGS1_WT-UTR, or DEGS1_edALL-UTR for d17-labeled ceramides at 2 hours and 24 hours after d17-sphinganine addition 2 days post-transfection. d17-sphinganine was added to both groups at the same time and collected at different times. The levels of C14, C16, C18, C22, C24:1, or C24 acyl chain length ceramides were normalized to total cell count. N=2 for 2 hr d17-sphinganine, edALL group; N=3 for all other groups. * denotes p < 0.05, ** denotes p < 0.01, *** denotes p < 0.001, **** denotes p < 0.0001 using two-way ANOVA with Tukey’s multiple comparisons test.

**Supplementary Figure 5.**
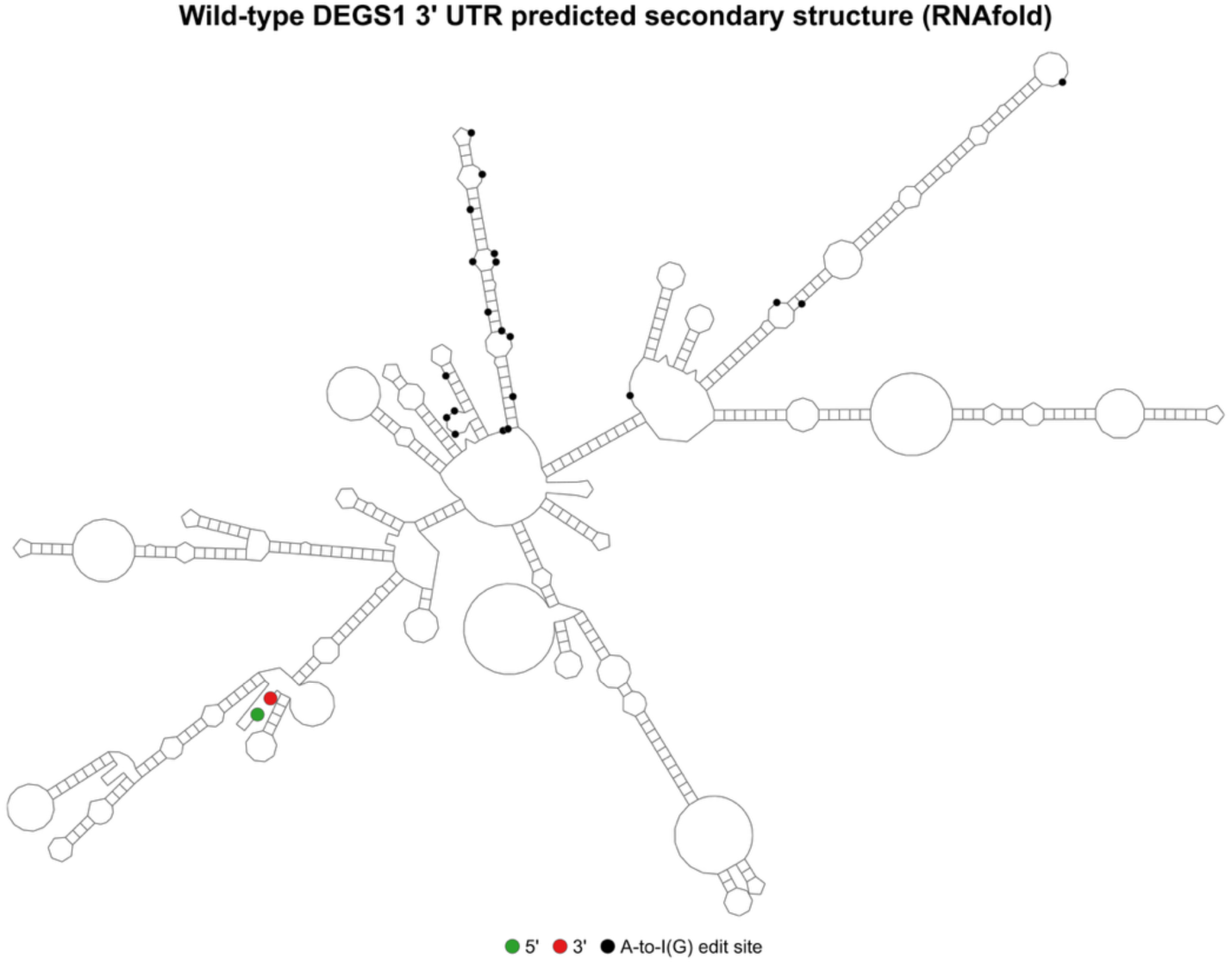
RNA edit sites mapped onto predicted secondary structure of DEGS1 3’ UTR. Secondary structure of the *DEGS1* 3’ UTR using RNAfold. Green and red dots represent 5’ and 3’ ends respectively. Black dots represent A-to-I(G) edit sites at specific stem and loop structures of the 3’ UTR.

**Supplementary Figure 6.**
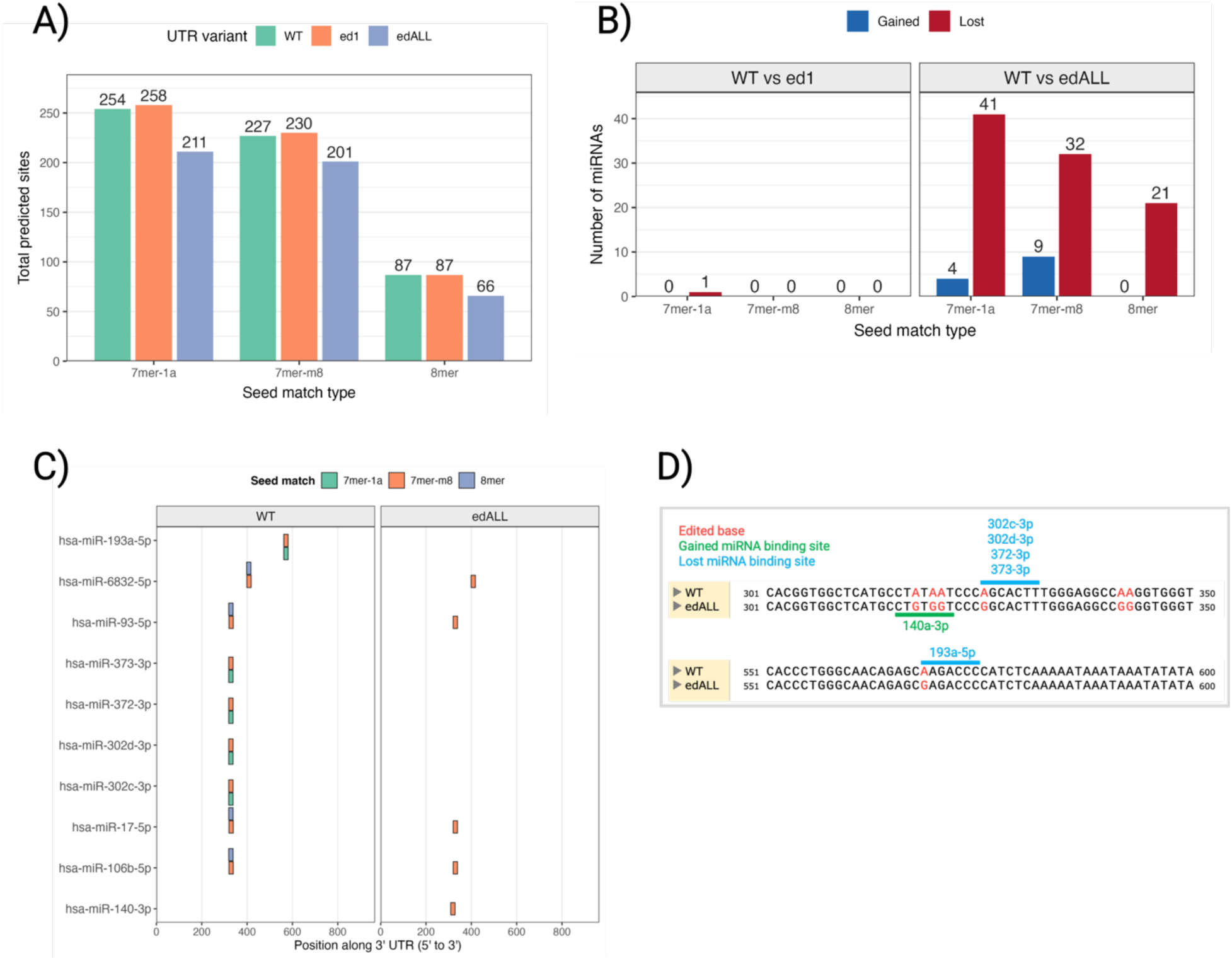
Gain and loss of miRNA binding sites upon RNA editing of DEGS1 3’ UTR. Changes in miRNA binding sites. **(A)** Predicted number of edit sites for each UTR variant for a given seed match type. **(B)** Comparison of unique miRNA binding sites lost or gained from ed1 and edALL compared to WT 3’ UTR. **(C)** miRNAs associated with ALS by cross-referencing miRNAs with HMDD and PubMed literature. miRNAs were mapped onto the location of the 3’ UTR. **(D)** Schematic showing lost or gained miRNAs associated with ALS literature mapped onto the 3’ UTR of DEGS1 when comparing edALL to WT.

## Supplementary File Legends

**Supplementary file 1. List of differentially edited sites**

All sites with significant differential editing (Benjamini–Hochberg FDR < 0.05).

**Supplementary file 2. Ingenuity Pathway Analysis canonical pathways by tissue**

Canonical-pathway enrichment of differentially edited genes (nominal P < 0.001 and absolute effect > 0.5). Table reports pathways reaching nominal significance (P < 0.05).

**Supplementary file 3. WT and edited DEGS1 3’ UTR sequences**

Nucleotide sequence of *DEGS1* WT, ed1 and edALL 3’ UTR variants used in this study

**Supplementary file 4. Summary of miRNA binding sites lost or gained and associated ALS literature**

Description of miRNA binding changes and its association with ALS.

