## Supplementary File 3 for "RNA Editing of DEGS1 Links Genetic Risk to Ceramide Dysregulation and Astrocyte Toxicity in ALS"

**DEGS1 3’ UTR sequences**

**DEGS1_WT-UTR:**

ATATCATTAGTGCCAAAGGGATTCTTCTCCAAAACTTTAGATGATAAAATGGAATTTTTGCATTATTAAACTTGAGACCAGTGATGCTCAGAAGCTCCCCTGGCACAATTTCAGAGTAAGAGCTCGGTGATACCAAGAAGTGAATCTGGCTTTTAAACAGTCAGCCTGACTCTGTACTGCTCAGTTTCACTCACAGGAAACTTGTGACTTGTGTATTATCGTCATTGAGGATGTTTCACTCATGTCTGTCATTTTATAAGCATATCATTTAAAAAGCTTCTAAAAAGCTATTTCGCCAGGCACGGTGGCTCATGCCTATAATCCCAGCACTTTGGGAGGCCAAGGTGGGTGGATCACCTGAGGTCAGGAGTTCGAGACCAGCCTGGCCAACACGGTGAAACCCCATCTCTACTAAAAATGCAAAAATTAGCCGGGCGTGGCGGCACATGCCTGTAATCCCAGCTACATGGGAGGCTGAGGTGGGAGAATTGCTTGAACCCAGGAGGCGGAGGCAGAGGCTGCAGTGACCCAAGATTGTGCCACTGCACTCCACCCTGGGCAACAGAGCAAGACCCCATCTCAAAAATAAATAAATATATATAAAAAATAAAAAGCTATTTCTAGTTTATTTCACTATAAAGTTTTGCTTTATTAAAAAGCTAATAAACAGCTATTAATCACAGTGTATTAGTATTTGTTACATTTTTGTATTTCACTATCTTTATACTATATAATATGGTAACTTGGGTACCGGGGGAACTTTAAAATTTCATCTCAAAAATAATTTTTAAAAAGCCTGAGGTATGATATAGCATAAAAGATTGAGATGAAAATATATTTCCCTGTAAGCTGAATTACTCATTTAAAAATTTTAACTTCTATATGGGACCCGAATTAGACACTGCTGAATCCTGTACAGCCTTACTCATAAATAAAGTACTTACTGAATTTCCACCATTCAAA

**DEGS1_ed1-UTR:**

ATATCATTAGTGCCAAAGGGATTCTTCTCCAAAACTTTAGATGATAAAATGGAATTTTTGCATTATTAAACTTGAGACCAGTGATGCTCAGAAGCTCCCCTGGCACAATTTCAGAGTAAGAGCTCGGTGATACCAAGAAGTGAATCTGGCTTTTAAACAGTCAGCCTGACTCTGTACTGCTCAGTTTCACTCACAGGAAACTTGTGACTTGTGTATTATCGTCATTGAGGATGTTTCACTCATGTCTGTCATTTTATAAGCATATCATTTAAAAAGCTTCTAAAAAGCTATTTCGCCAGGCACGGTGGCTCATGCCTGTAATCCCAGCACTTTGGGAGGCCAAGGTGGGTGGATCACCTGAGGTCAGGAGTTCGAGACCAGCCTGGCCAACACGGTGAAACCCCATCTCTACTAAAAATGCAAAAATTAGCCGGGCGTGGCGGCACATGCCTGTAATCCCAGCTACATGGGAGGCTGAGGTGGGAGAATTGCTTGAACCCAGGAGGCGGAGGCAGAGGCTGCAGTGACCCAAGATTGTGCCACTGCACTCCACCCTGGGCAACAGAGCAAGACCCCATCTCAAAAATAAATAAATATATATAAAAAATAAAAAGCTATTTCTAGTTTATTTCACTATAAAGTTTTGCTTTATTAAAAAGCTAATAAACAGCTATTAATCACAGTGTATTAGTATTTGTTACATTTTTGTATTTCACTATCTTTATACTATATAATATGGTAACTTGGGTACCGGGGGAACTTTAAAATTTCATCTCAAAAATAATTTTTAAAAAGCCTGAGGTATGATATAGCATAAAAGATTGAGATGAAAATATATTTCCCTGTAAGCTGAATTACTCATTTAAAAATTTTAACTTCTATATGGGACCCGAATTAGACACTGCTGAATCCTGTACAGCCTTACTCATAAATAAAGTACTTACTGAATTTCCACCATTCAAA

**DEGS1_edALL-UTR:**

ATATCATTAGTGCCAAAGGGATTCTTCTCCAAAACTTTAGATGATAAAATGGAATTTTTGCATTATTAAACTTGAGACCAGTGATGCTCAGAAGCTCCCCTGGCACAATTTCAGAGTAAGAGCTCGGTGATACCAAGAAGTGAATCTGGCTTTTAAACAGTCAGCCTGACTCTGTACTGCTCAGTTTCACTCACAGGAAACTTGTGACTTGTGTATTATCGTCATTGAGGATGTTTCACTCATGTCTGTCATTTTATAAGCATATCATTTAAAAAGCTTCTAAAAAGCTATTTCGCCAGGCACGGTGGCTCATGCCTGTGGTCCCGGCACTTTGGGAGGCCGGGGTGGGTGGATCGCCTGGGGTCGGGAGTTCGGGACCGGCCTGGCCGGCACGGTGGGACCCCGTCTCTACTAAAAATGCAAAAATTGGCCGGGCGTGGCGGCACATGCCTGTAATCCCAGCTACATGGGAGGCTGAGGTGGGAGAGTTGCTTGAACCCAGGAGGCGGAGGCAGAGGCTGCAGTGACCCAGGATTGTGCCACTGCACTCCACCCTGGGCAACAGAGCGAGACCCCATCTCAAAAATAAATAAATATATATAAAAAATAAAAAGCTATTTCTAGTTTATTTCACTATAAAGTTTTGCTTTATTAAAAAGCTAATAAACAGCTATTAATCACAGTGTATTAGTATTTGTTACATTTTTGTATTTCACTATCTTTATACTATATAATATGGTAACTTGGGTACCGGGGGAACTTTAAAATTTCATCTCAAAAATAATTTTTAAAAAGCCTGAGGTATGATATAGCATAAAAGATTGAGATGAAAATATATTTCCCTGTAAGCTGAATTACTCATTTAAAAATTTTAACTTCTATATGGGACCCGAATTAGACACTGCTGAATCCTGTACAGCCTTACTCATAAATAAAGTACTTACTGAATTTCCACCATTCAAA
